# SMCHD1 variants may induce variegated expression in Facio Scapulo Humeral Dystophy and Bosma Arhinia and microphtalmia syndrome

**DOI:** 10.1101/2021.05.17.444338

**Authors:** Camille Laberthonnière, Raphaël Chevalier, Camille Dion, Mégane Delourme, David Hirst, José Adélaïde, Max Chaffanet, Shifeng Xue, Karine Nguyen, Bruno Reversade, Jérôme Déjardin, Anais Baudot, Jérôme D. Robin, Frédérique Magdinier

## Abstract

An expanding number of genetic syndromes are linked to mutations in genes encoding factors that guide chromatin organization. Recently, distinct genetic syndromes have been linked to mutations in the *SMCHD1* gene. However, the function of this non-canonical SMC protein remains partly defined in Human tissues. To address this question, we determined its epi-signature in type 2 Facio Scapulo Humeral Dystrophy (FSHD2) and Bosma Arhinia and Microphtalmia Syndrome (BAMS) linked to heterozygous mutations in this gene. By combining RNA-Seq, DNA methylation profiling and ChIP-Seq, we showed that SMCHD1 regulates repressed chromatin but also *cis*-regulatory elements and enhancers. Our results emphasize dual functions for SMCHD1, in chromatin compaction, chromatin insulation and gene regulation with variable outcomes and targets depending on tissues. We propose that altered DNA methylation and long-range chromatin organization at a number of loci required for development and tissue differentiation, trigger variegated gene expression in rare genetic diseases linked to heterozygous *SMCHD1* mutations.

## Introduction

Based on the presence of an SMC hinge domain, the SMCHD1 (Structural Maintenance of Chromosomes flexible Hinge Domain containing 1) chromatin associated factor belongs to the SMC family of chromosomal proteins. However, this factor that forms homodimers through interaction between its SMC hinge domains does not participate in the tripartite ring complex formed by other Cohesins (Chen et al., 2016). In the mouse, Smchd1 has been mainly associated with X inactivation since its depletion causes embryonic lethality with absence of repression of genes on the inactive X chromosome (Blewitt et al., 2008; Gendrel et al., 2012; Gendrel et al., 2013). Besides, the protein is implicated in the topological conformation of the inactive X chromosome (Gdula et al., 2019; Jansz et al., 2018; Wang et al., 2018) and formation A and B mega-domains separated by the Dxz4 macrosatellite (Darrow et al., 2016; Deng et al., 2015; Giorgetti et al., 2016; Wang et al., 2018). However, the precise mechanism, likely dependent on Xist or H3K27me3 enrichment, underlying this process remains partially understood. Smchd1 is also implicated in the regulation of heterochromatin, repetitive DNA sequences or clustered imprinted genes and monoallelically expressed Protocadherin genes (Blewitt et al., 2008; Blewitt et al., 2005; Brideau et al., 2015; Gendrel et al., 2013; Mason et al., 2017; Massah et al., 2014; Mould et al., 2013; Nozawa et al., 2013; Wanigasuriya et al., 2020). SMCHD1 has also been found at telomeres with a direct correlation between telomere length and SMCHD1 enrichment (Dejardin and Kingston, 2009; Grolimund et al., 2013). Heterozygous germline *SMCHD1* mutations have been reported in at least three distinct rare human genetic diseases, type 2 Facio-Scapulo-Humeral muscular dystrophy (FSHD2) (Gaillard et al., 2016; Lemmers et al., 2012; Sacconi et al., 2012), Bosma Arhinia and Microphtalmia Syndrome (BAMS) (Gordon et al., 2017; Shaw et al., 2017) and Isolated Hypogonadotrophic Hypogonadism (IHH) with Combined Pituitary Hormone Deficiency (CHPD) and Septo-Optic Dysplasia (SOD) (Kinjo et al., 2020). The gene is also often deleted in patients carrying an 18p deletion and presenting symptoms that differ from the three other diseases (Cody et al., 2015; Hasi-Zogaj et al., 2015) or a typical FSHD phenotype (Lemmers et al., 2015).

FSHD is an autosomal dominant muscular dystrophy, linked in 95% of cases to shortening of an array of 3.3 kb macrosatellite elements (D4Z4) in the distal region of the 4q arm (Wijmenga et al., 1992). In 2-3 % of FSHD cases, the *D4Z4* array is not shortened but patients carry a mutation in *SMCHD1* (FSHD2; OMIM 158901) (Lemmers et al., 2012). *D4Z4* is GC-rich (70%) (Lyle et al., 1995) and hypomethylated in FSHD1 and FSHD2 patients, as a consequence of D4Z4 array shortening or *SMCHD1* mutation respectively (Gaillard et al., 2014; Hartweck et al., 2013; Roche et al., 2019; van Overveld et al., 2003).

BAMS (OMIM 603457) is an extremely rare and striking condition characterized by complete absence of a nose with or without ocular defects but no sign of muscular dystrophy (Gordon et al., 2017; Shaw et al., 2017) likely resulting from a defect of the nasal placodes or surrounding neural crest-derived tissues migration.

In FSHD, *SMCHD1* mutations are dispersed across the whole coding sequence while in BAMS, mutations are clustered within exons 3 to 13, spanning a GHKL-type ATPase domain and an associated region immediately C terminal to it (Gordon et al., 2017; Shaw et al., 2017). BAMS has been linked to a gain of function of the SMCHD1 ATPase domain (Gordon et al., 2017; Gurzau et al., 2018; Shaw et al., 2017) while in FSHD2 variants are mostly missense, splice and truncating mutations likely leading to a loss of function or haploinsufficiency (Dion et al., 2019; Gordon et al., 2017; Gurzau et al., 2018; Lemmers et al., 2012). D4Z4 hypomethylation is a feature of FSHD (de Greef et al., 2010; de Greef et al., 2009; Sacconi et al., 2012; van Overveld et al., 2005; van Overveld et al., 2003) but the repetitive macrosatellite is also hypomethylated In BAMS, as well as in 18p11.32 hemizygous patients carrying a deletion of the *SMCHD1* locus (Dion et al., 2019; Gordon et al., 2017; Shaw et al., 2017). Thus, SMCHD1 haploinsufficiency, loss or gain of function mutations are all associated with hypomethylation and chromatin relaxation of the D4Z4 macrosatellite with however different phenotypical consequences.

*SMCHD1* mutations in BAMS or FSHD2 are not associated with the X inactivation defects described in mice (Dion et al., 2019). Human mutations in zebrafish and mouse models do not recapitulate the BAMS or FSHD phenotype (Gordon et al., 2017; Shaw et al., 2017). For a number of loci such as *HOX* genes, imprinted or *PCDH* loci, SMCHD1 function might show some commonalities between mouse and human tissues (Gendrel et al., 2013; Jansz et al., 2018; Mason et al., 2017; Massah et al., 2014; Mould et al., 2013). However, the peculiarity of the human SMCHD1 during cell fate transitions remains to be established.

Taking advantage of samples from patients affected either with BAMS or FSHD2, we investigated the implication of this protein in chromatin regulation in human cells at the genome-wide scale. We uncovered pleiotropic roles for this protein in heterochromatinization but also in insulating against repressive chromatin marks or in the chromatin conformation of enhancer elements. Overall, SMCHD1 connects two major determinants of genome regulation, DNA methylation and chromatin architecture with variable outcomes depending on the (epi)genomic context or tissue. Given its potential implication in the regulation of genes entailed for development and tissue differentiation, we further propose that variegated gene expression triggered by SMCHD1 partial deficiency might contribute to the clinical spectrum of SMCHDnopathies.

## Results

### Mutations in *SMCHD1* are associated with changes in the expression of a small subset of protein coding genes

Given the absence of a clear function for SMCHD1 in the regulation of the human genome but its implication in several rare genetic diseases, we first seek at identifying genes dysregulated in primary cells from patients carrying a mutation in *SMCHD1* and affected with either BAMS or FSHD2 (Figure 1A, Table S1). To this aim, we performed RNA-Seq analysis in primary fibroblasts from three patients for each disease. In comparison to healthy donors (controls), we found 1507 differentially expressed genes (DEGs) in BAMS and 1134 in FSHD2 cells with an overlap of 626 genes (Figure 1B, Figure S1A, B; fold change ≥ 2; p value < 0.05) between the two groups of patients. Only a few genes show an opposite trend of expression (up in BAMS and down in FSHD2: *KRT7, HAPLN1* or vice versa, *EMB, MEOX2* and *LCNL1*). Analysis of biological pathways (BP) revealed enrichment in gene ontology (GO) terms corresponding to factors involved in development and patterning with 11 out of the 15 most conserved pathways shared between the two diseases (figure 1C).

**Figure 1.**
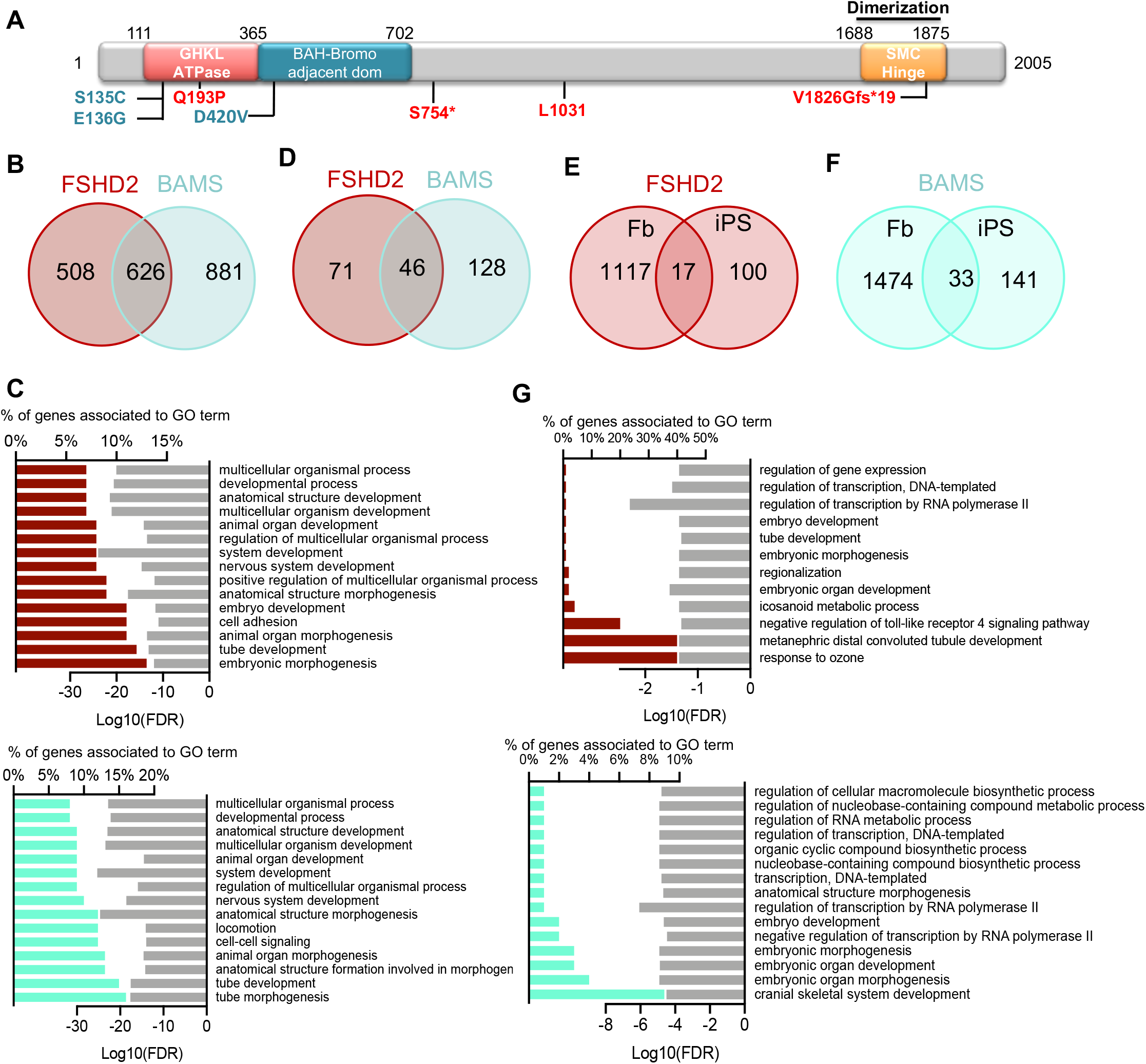
Gene expression profiling in fibroblasts and induced pluripotent stem cells from patients affected with BAMS or FSHD2. **A**. Schematic representation of the SMCHD1 protein with position of the different mutations in BAMS (cyan) or FSHD2 (red) patients. **B**. Venn diagrams for comparison of genes that are differentially expressed in FSHD2 and BAMS primary fibroblasts compared to controls with a fold-change >2 and <-2 and an FDR <0.05. **C**. Gene Ontology (GO) for Biological pathways (BP) corresponding to enrichment analysis of DEGs in FSHD2 (red, upper panel) or BAMS (cyan, lower panel) *vs* Control fibroblasts filtered on -2>FC>2 and FDR <0.05. Bar plots in the left represents the percentage of genes that are differentially expressed and associated with a GO-term shown in the right column. Light grey bars in the right represent the enrichment score (Log10 of False Discovery Rate) for each BP. **D**. Venn diagrams for comparison of genes that are differentially expressed in FSHD2 and BAMS induced pluripotent stem cells (hiPSCs) compared to controls. **E**. Venn diagrams for comparison of genes that are differentially expressed between FSHD2 fibroblasts and hiPSCs. **F**. Venn diagrams for comparison of genes that are differentially expressed between BAMS fibroblasts and hiPSCs. **G**. Gene Ontology for Biological pathways corresponding to enrichment analysis of DEGs in FSHD2 (red, upper panel) or BAMS (cyan, lower panel) *vs* Control hiPSCs filtered on -2>FC>2 and FDR <0.05. Bar plots in the left represents the percentage of DEGs and associated with GO-terms in the right column. Light grey bars in the right represent the enrichment score (Log10 of False Discovery Rate) for each GO-term.

Given the role for SMCHD1 chromatin regulation during pluripotency (Dion et al., 2019), we next asked whether mutations in this gene modulate gene expression after reprogramming. In induced pluripotent stem cells (hiPSCs) from patient’s and control’s fibroblasts (Table S1), we identified 174 DEGs in BAMS and 117 in FSHD2, with an overlap of 46 genes between the two diseases (Figure 1D; Figure S1C, D). Thus, after reprogramming, the expression of approximately 90% of the DEGs found in fibroblasts is reset with an overlap of 17 DEGs for FSHD2 (Figure 1E) and 33 genes in BAMS (Figure 1F). BPs correspond to various cellular functions or developmental processes with 3 out of 15 pathways common between the two diseases (Figure 1G).

In FSHD, SMCHD1 loss-of-function, D4Z4 chromatin relaxation and subsequent *DUX4* activation is proposed as the disease driver mechanism. We thus searched for DUX4 and its 422 DUX4 target genes (Geng et al., 2012) among DEGs. DUX4 is undetectable by RNA-Seq due to its low expression level and only a small number (8%) of DUX4 targets are among DEGs. Notably in fibroblasts, we found 19 genes in BAMS and 16 in FSHD2 including 5 common to both diseases (Figure S1E). In hiPSCs, only one gene (*OAS1*) was identified among DEGs in BAMS (Figure S1F).

Thus, fibroblasts with mutations in *SMCHD1* (BAMS or FSHD2) display differential expression of a number of genes associated with development and cell differentiation but are not distinguishable based on the expression profile of DUX4 or its target genes.

### Comparative methylation landscape between BAMS and FSHD2 primary cells

Considering the role of SMCHD1 in DNA methylation, we performed methylation profiling of primary fibroblasts (BAMS, n=3, FSHD2, n=3, controls, n=2) at early passages (<10) after sodium bisulfite conversion using the Illumina Infinium HumanMethylation 850 BeadChip (Moran et al., 2016). For each probe, the methylation level of CpGs covered by the array was determined by calculating the median DNA methylation β-values and standard deviation (SD) within samples groups to detect differential methylation with a 99% confidence. Probes with an absolute difference (mean beta values of 0.2) were considered as differentially methylated (DMP, Differentially Methylated Probe) (Bibikova et al., 2011). After batch effect correction and removal of probes corresponding to SNPs and sex chromosomes, density plot analyses and unsupervised hierarchical clustering indicate an overall overlap of profiles between samples (Figure S2A-D). We identified 29528 DMPs in BAMS (3% of probes) and 53709 DMPs in FSHD2 (6%) compared to controls including 12054 DMPs (20% of disease-associated DMPs) shared between the two diseases. Methylation is globally increased in BAMS cells but decreased in FSHD2 compared to controls (Figure 2A, Figure S3). The disparity between samples might be explained by the discrepancy in the proportion of HyperM probes in BAMS but the balanced proportion of HyperM and HypoM probes between FSHD2 samples (Figure 2B, C). Overall, in the two diseases, the distribution of DMPs by CpG content, categorized using the Illumina annotations is equivalent with a majority of DMPs corresponding to open seas (figure 2D-F), gene bodies and Internal Genomic Regions (IGR) (figure 2G-I).

**Figure 2.**
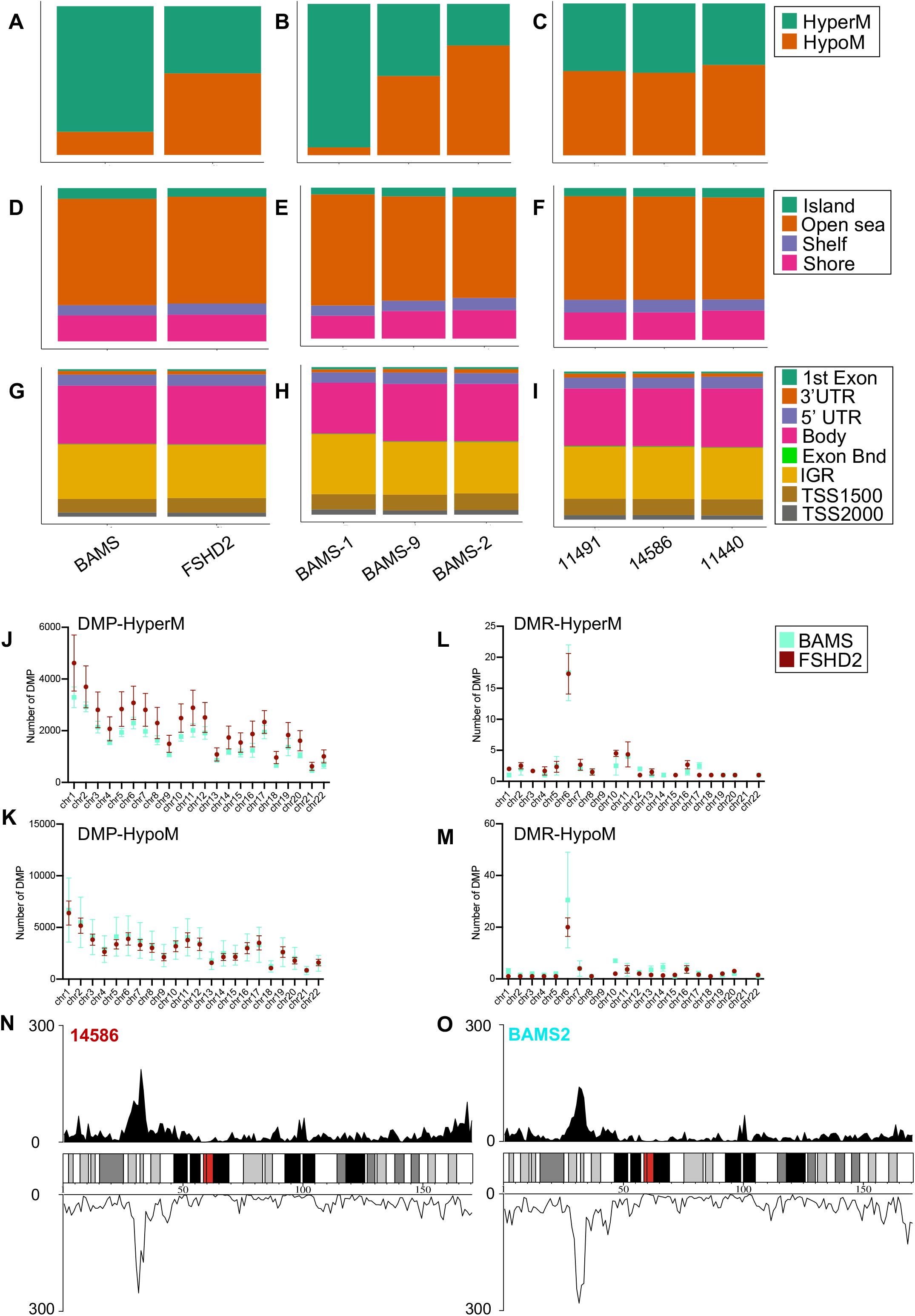
Distribution of DMPs relative to CpG content. **A-C**. Cumulative histograms of the distribution of hypo- and hypermethylated probes in BAMS and FSHD2 fibroblasts (**A**), individual BAMS samples (**B**) or individual FSHD2 samples (**C**). **D-F**. Cumulative histograms for DMP by CpG content relative to CpG islands, shores (2 kb flanking CpG islands), shelves (2 kb extending from shores) or open seas (isolated CpG in the rest of the genome) in BAMS and FSHD2 fibroblasts (**D**), individual BAMS samples (**E**) or individual FSHD2 samples (**F**). **G-I**. Cumulative histograms for DMP by features corresponding to genes first exon, 3’ UTR, 5’UTR, gene bodies, Exon boundaries, Internal genomic regions (IGR), probes located 1500 bp from transcription start sites (TSS1500) or 2000 bp from transcription start sites (TSS200), in BAMS and FSHD2 fibroblasts (**G**), individual BAMS samples (**H**) or individual FSHD2 samples (**I**). **J-K**. Distribution of hyperM (**J**) or hypoM (**K**) DMP relative to autosomes. **L-M**. Distribution of hyperM (**L**) or hypoM (**M**) DMRs relative to autosomes for FSHD2 (red) or BAMS (cyan) primary fibroblasts. **N-O**. Representative distribution of the DNA methylation profile of DMR on chromosome 6. DMPs were represented as a density measured in bins Bin size chr6 = 1 000 000 bp. Upper graph corresponds to hyperM DMR, lower graph to hypoM DMRs in FSHD2-14586 (**N**) or BAMS2 (**O**) cells relative to healthy controls.

Next, we used the probe lasso method (Butcher and Beck, 2015) to identify differentially methylated regions (DMRs) in fibroblasts. When compared to controls, we found 70 to 263 DMRs in BAMS and 49 to 110 DMRs in FSHD2, distributed across autosomes (Figure 2J-M). Most DMRs are located in non-coding regions, *i*.*e* open seas, shores, bodies or IGRs. In the top DMRs, several map to chromosome 6 (Figure 2N-O, Figure S4), in particular at the 6p21.32-6p22.1 band encompassing the Major Histocompatibility Complex (MHC) locus and characterized by a high proportion of genes encoding chromatin factors and micro RNAs. Using stringent parameters, we then selected DMRs shared between BAMS and FSHD2 and sorted HyperM and HypoM DMRs as well as DMRs with opposite methylation status. The ChromHMM track from NHEK cells (normal epidermal keratinocytes, Broad Institute; Encode) was used to predict association to functional elements. A large proportion of these DMRs corresponds to repressed chromatin/heterochromatin or repressed and poised promoters (Figure 3A,B). Remarkably, we also observed a high proportion of HypoM or HyperM DMRs that correspond to enhancers. We concluded on a dual impact of *SMCHD1* mutation triggering either unwanted DNA methylation or protecting against this epigenetic modification (Figure 3D). BPs associated with genes in the vicinity of these DMR encode factors involved in developmental processes, including skeletal system development (Figure 3E).

**Figure 3.**
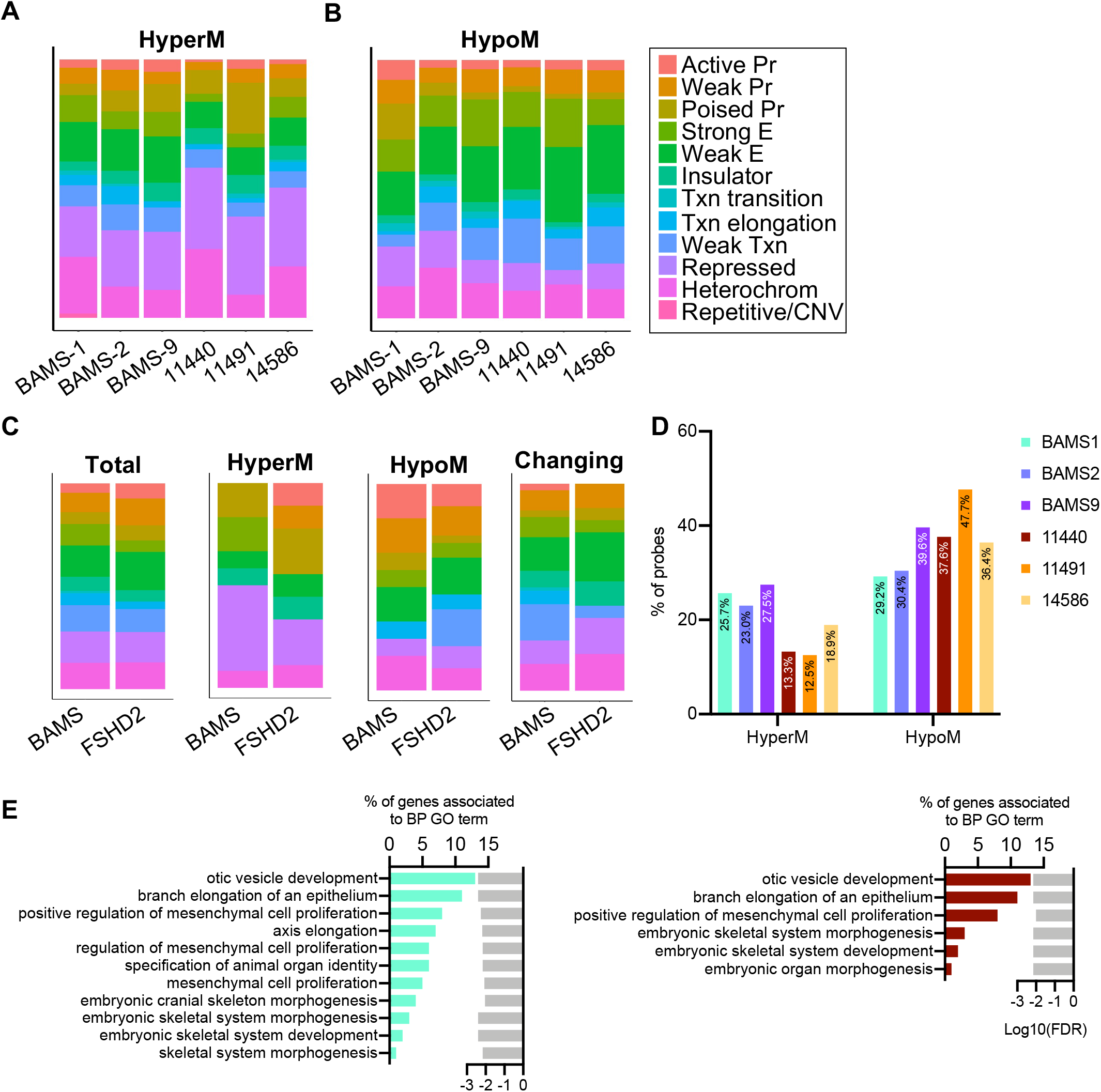
Profiling of DNA methylation in SMCHD1 deficient cells. **A-B**. DMRs with an FDR adjusted pvalue < 0.05 were analyzed for chromHMM features using NHEK cell annotations and ploted using R package ggplot2 (v3.3.3). Cumulative histograms for HyperM (**A**) or HypoM (**B**) probes are presented. **C**. From left to right, barplots of DMRs shared between patient when compared to controls (total); shared hypermethylated DMRs (HyperM); shared hypomethylated DMRs (HypoM); DMRs with a different methylation profile between patients (Changing). **D**. Graph displaying the percentage of hypermethylated or hypomethylated probes at enhancers in the different BAMS and FSHD2 samples. **E**. Biological pathways overrepresented for BAMS (cyan, left) or FSHD2 (red, right) DMRs. Bar plots in the left represent the percentage of genes and associated with GO-terms listed in the right column. Light grey bars in the right represent the enrichment score (Log10 of False Discovery Rate) for each GO-term.

In SMCHD1-deficient cells, the high proportion of HypoM probes confirms a role for this factor in chromatin compaction. Increased methylation of heterochromatin also indicates that SMCHD1 might shield against heterochromatinization. Hypo- or Hyper methylation of enhancers uncovers a role for SMCHD1 in the regulation of *cis*-regulatory elements in human tissues. We concluded that SMCHD1 deficiency might promote the methylation-dependent ectopic activation or repression of *cis*-regulatory elements in patient’s cells, with variable repercussions depending on the type of mutation.

### SMCHD1 regulates DNA methylation of a small subset of sequences in pluripotent cells

Next, we evaluated the methylation landscape after cell reprogramming of control and SMCHD1-deficient cells. We observed a global reset in the methylation profile with only 4-6 DMRs in BAMS hiPSCs and 3 DMRs in FSHD2 hiPSCs (Figure S5A, B). We observed an opposite profile compared to fibroblasts with a majority of HypoM probes in BAMS but a majority of HyperM probes in FSHD2 (Figure S5C, D). The distribution of DMPs differs between the two diseases with a higher proportion of DMP in CpG islands in BAMS and a higher proportion in open seas and shelves for FSHD2 hiPSCs (Figure S5D). As in fibroblasts, most DMPs are located in gene bodies and IGRs (Figure S5E).

Overall, this indicates a massive reset in the DNA methylation profile after cell reprogramming, with variable impacts depending on the type of mutation in the distribution of methylated CGs, as reported for the D4Z4 macrosatellite (Dion et al., 2019).

### Mutations in *SMCHD1* are associated with methylation changes at *HOX* gene loci

In the mouse, Smchd1 regulates large gene clusters such as *Hox* loci (Jansz et al., 2018). We then investigated changes in DNA methylation at the four human *HOX* genes clusters and compared their methylation level to *HOX* genes expression.

We identified DMRs with features of Polycomb repressed chromatin (*HOXA13/HOTTIP*), poised promoter (*HOXB2, HOXB6)* or enhancer (*HOXC4; C5; C6)* (Figure 4A-D; Figure S6 A-D). As all CGs are not covered by the Epic Array, we first performed in depth methylation profiling of these DMRs by sodium bisulfite sequencing (BSS) in fibroblasts and after reprogramming. We noted variable profiles in fibroblasts and hiPSCs. *HOXA13* methylation is identical between fibroblasts and hiPSCs in controls. This DMR is markedly hypomethylated in one of the FSHD2 samples compared to the other FSHD2 and BAMS samples with no apparent correlation for mutations altering the ATPase activity. In hiPSC, the opposite is observed with an increased methylation in cells derived from hypomethylated fibroblasts (14586) and an absence of methylation when the DMR is methylated in fibroblasts (120521C, BAMS1-9) (Figure S7A). *HOXB2* DMR is unmethylated in the majority of samples (Figure S7B). *HOXB6*, methylation is low except in BAMS-2 fibroblasts (gain of SMCHD1 ATPase activity (Gurzau et al., 2018)) but increases after reprogramming in one of the FSHD2 samples (14586; loss of ATPase activity (Dion et al., 2019)) (Figure S7C). Interestingly, for *HOXC4*, we observed an absence of methylation in samples carrying a mutation affecting the ATPase activity (FSHD2-14586, BAMS-1, BAMS-2) and an increased methylation in other patient’s fibroblasts (120521C, BAMS-9). Demethylation of this DMR occurs in all samples after reprogramming (Figure S7C). Twenty-two *HOX* genes are differentially expressed in BAMS and FSHD2 fibroblasts without any obvious correlation between gene expression and methylation changes (Figure 4E, F). Thus, as observed in the mouse and besides its role in heterochromatin condensation SMCHD1 contributes to the regulation of DNA methylation of autosomal loci with variable impact of *SMCHD1* germline mutations on their methylation profile as observed for D4Z4 (Dion et al., 2019).

**Figure 4.**
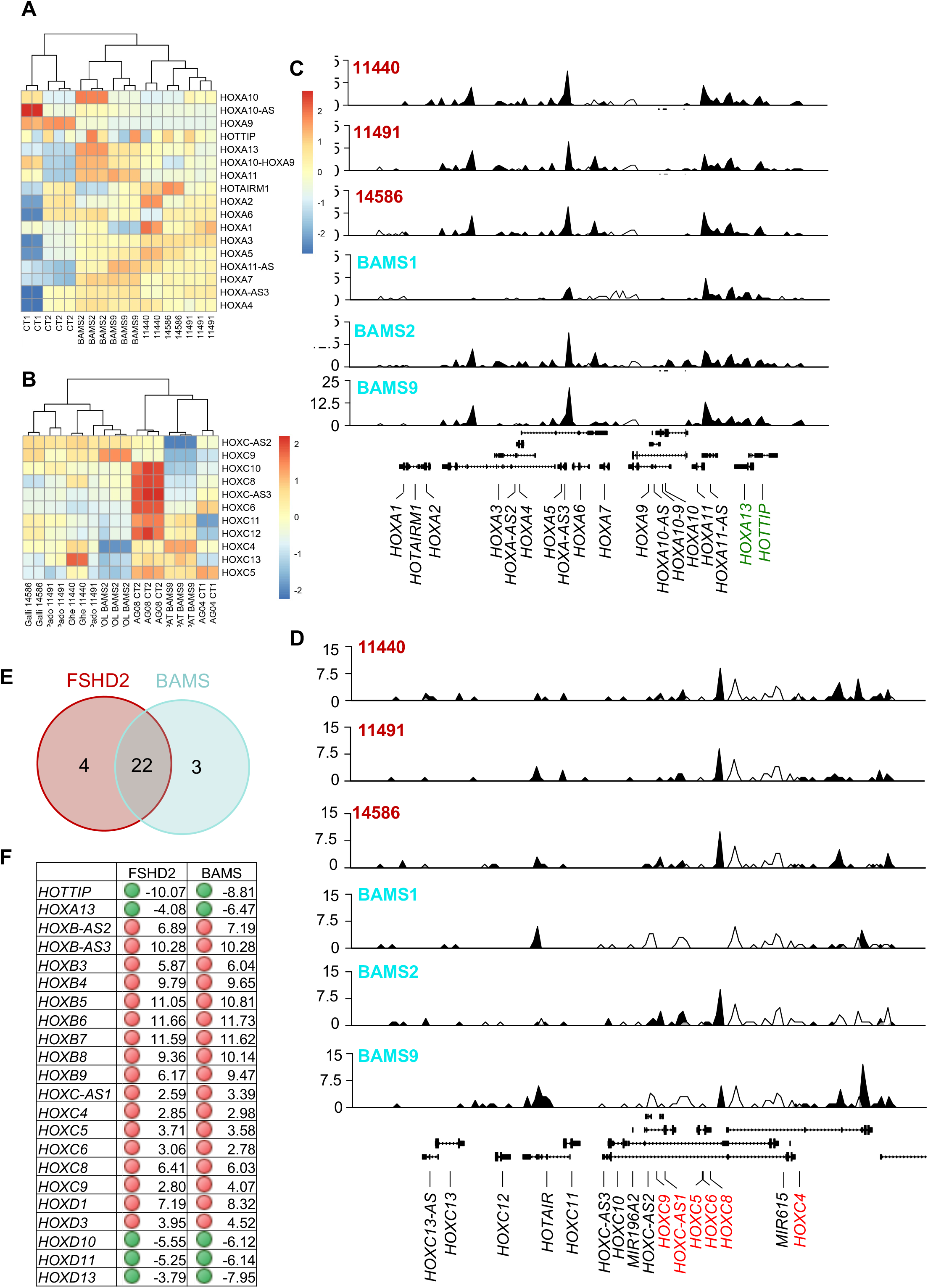
Germline *SMCHD1* mutations result in changes in *HOX* genes expression and methylation profiles. **A**. Hierarchical clustering of differential methylation at *HOXA* gene loci (chr7: 27,064,176-27,227,600) in BAMS, FSHD2 and Controls with Ward’s method on Euclidian distance for the different samples. **B**. Hierarchical clustering of differential methylation at *HOXC* gene loci in BAMS, FSHD2 and Controls with Ward’s method on Euclidian distance for the different samples. **C**. Representative distribution of the DNA methylation profile of DMR at the *HOXA* locus in the different FSHD2 (11440, 11491, 14586, red) or BAMS (BAMS1, BAMS2, BAMS9) samples. Black curves correspond to HyperM probes, white curves to HypoM ones. **D**. Representative distribution of the DNA methylation profile of DMR at the *HOXC* locus (chr12: 53,908,560-54,069,307) in the different FSHD2 (11440, 11491, 14586, red) or BAMS (BAMS1, BAMS2, BAMS9) samples. **E**. Venn diagrams for comparison of *HOX* genes that are differentially expressed in FSHD2 and BAMS fibroblasts compared to controls with a fold-change >2 and <-2 and an FDR <0.05. **F**. Table of differentially expressed *HOX* genes in FSHD2 and BAMS cells. Red dots correspond to genes that are upregulated and green dots, to downregulated ones. For each gene, Fold Change (FC) is indicated.

### Somatic SMCHD1 invalidation does not modify DNA methylation

D4Z4 hypomethylation is a hallmark of patients carrying a germline *SMCHD1* mutation (Dion et al., 2019; Gordon et al., 2017; Hartweck et al., 2013; Shaw et al., 2017). Conversely, somatic *SMCHD1* invalidation does not cause D4Z4 hypomethylation and SMCHD1 binding to the macrosatellite is DNA methylation-independent (Dion et al., 2019). To address whether these observations also apply to other sequences, we analyzed the four *HOX* DMRs in HEK and HEK *SMCHD1* KO cells. SMCHD1 binds to the different DMRs (Figure S8A) but somatic invalidation of this protein does not cause any change in DNA methylation (Figure S8B). Absence of SMCHD1 does not impact *HOXB* genes expression but is associated with an increased expression of genes controlled by the *HOXA13* and *HOXC4/5/6* DMRs (Figure S8C) highlighting a possible role for SMCHD1 in the regulation of a number of *HOX* genes. We confirmed the absence of effect of *SMCHD1* somatic invalidation on DNA methylation *in vivo*, by analyzing D4Z4 methylation in 30 breast cancer specimen with a loss of heterozygosity (LOH) at the 18p locus encompassing *SMCHD1*. As in HEK KO cells, D4Z4 is not significantly hypomethylated (Figure S9) confirming that this protein is dispensable for DNA methylation maintenance *in vitro* and *in vivo*.

### SMCHD1 deficiency leads to acute loss of H3K27me3 at enhancers in skeletal muscle cells

Our genome-wide methylation analysis revealed a role for SMCHD1 in the regulation of enhancer elements, associated to genes involved in development and cell differentiation. Thus, we next sought to identify gene regulatory circuits that depend on SMCHD1 in tissue affected in the disease and define their associated chromatin features. To this aim, we differentiated hiPSC into skeletal muscle fibers and mapped the activating H3K4me3 and repressive H3K27me3 marks together with distribution of the CTCF insulating factor by ChIP-seq. Peaks are distributed along the 22 autosomes. Compared to the methylation profile in fibroblasts, we did not observe any disproportional enrichment at chromosome 6 or any other autosome (Figure S10). The three categories of samples (Control, BAMS and FSHD2) display the same proportion of H3K4me3 peaks but a decrease in CTCF and H3K27me3 peaks in BAMS and FSHD2 (Figure S10) together with variable peaks width (Figure S10, p value <0.0001).

CTCF, H3K4me3 or H3K27me3 enrichment relative to chromatin features was determined using the ChromHMM track for Human Skeletal Muscle Myoblasts (HSMM) annotations (Figure 5A). Regarding H3K4me3, we did not observe any major change between conditions except a modest increase in the proportion of active promoters enriched in H3K4me3 in BAMS (38%) and FSHD2 (36.5%) cells compared to controls (33%) (Figure 5A). CTCF peaks-associated features are evenly distributed between samples, with a slight decrease at active promoters in BAMS and at heterochromatin in FSHD2.

**Figure 5.**
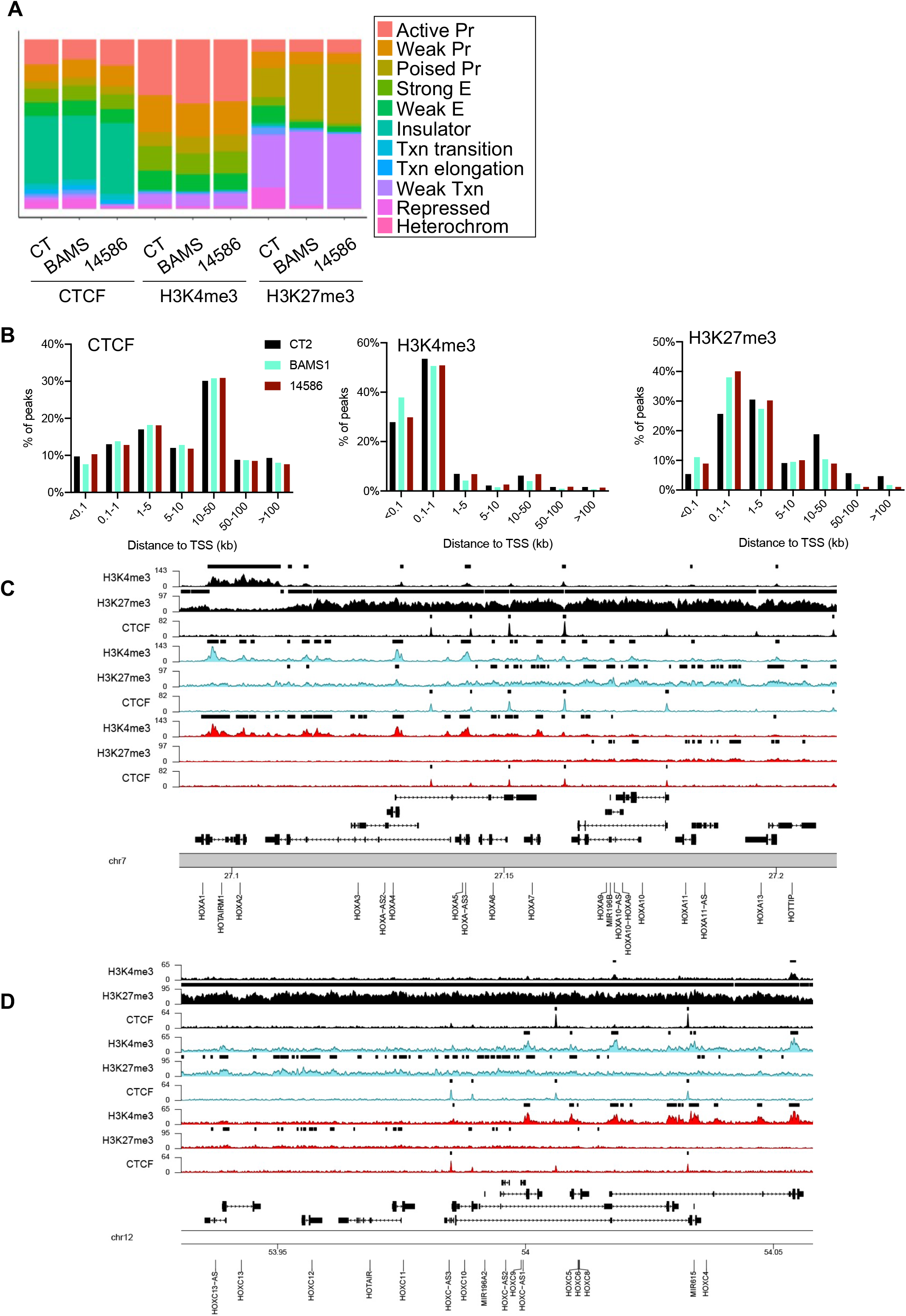
Chromatin profiling of hiPSC-derived muscle fibers from SMCHD1-deficient patients. **A**. Distribution of peaks enriched in CTCF, H3K4me3 or H3K27me3 relative to chromatin features determined using the ChromHMM track in muscle fibers derived from Control, BAMS1 or FSHD2 (14586) hiPSCs. Peaks with at least a qvalue < 0.05 in one replicate were analyzed for chromHMM features using HSMM (Human Skeletal Muscle Myoblasts) cells annotations and represented as barplot using R package ggplot2 (v3.3.3). **B**. Distribution of distances from Transcription Start Sites (TSS) for peaks enriched in CTCF, H3K4me3 or H3K27me3 in Controls (CT), BAMS or FSHD2 cells. Peaks distribution to TSS was assessed using R package chipenrich (v2.14.0). **C**. Profiling of the *HOXA* genes locus (chr7: 27,090,339-27,211,142, 120803 bp) in the different samples (Controls, black; BAMS, cyan or FSHD2, red) for H3K4me3, H3K27me3 or CTCF enrichment. **D**. Profiling of the *HOXC* genes locus (chr12: 53,930,562-54,058,000; 127438 bp) in the different samples (Controls, black; BAMS, cyan or FSHD2, red) for H3K4me3, H3K27me3 or CTCF enrichment.

Most changes concern the distribution of H3K27me3. Enrichment is decreased at heterochromatin (12.1% in Controls, 1.9% in BAMS and 0.72% in FSHD2) consistent with a role for SMCHD1 in chromatin compaction. However, we noted an increase of this mark at repressed chromatin (31.1% in Controls, 43.3% in BAMS, 43.9% in FSHD2) and poised promoters in BAMS (32.2%) and FSHD2 (35.7%) cells *vs* controls (17.3%) suggesting that SMCHD1 might shield against the deposition of repressive chromatin marks.

The most remarkable change occurs at enhancers with a deprivation in H3K27me3 in diseases (BAMS: 5%; FSHD2: 3.75%) compared to controls (15.1%) that might correlate with the slight increase in H3K4me3-enriched active promoters.

Relative to transcription start sites (TSS), we observed a similar distribution for CTCF and H3K4me3 peaks between samples but an increase in H3K27me3 at a short distance from TSS (0.1-1kb) in patient’s samples (Figure 5B) that correlates with the increased number of poised promoters harboring this mark (Figure 5A, Table S6) in BAMS and FSHD2 cells. Notably, depletion in H3K27me3 at distance from TSS (10 to >100 kb, Figure 5B) correlates with the drastic decrease in enhancers harboring this mark in BAMS and FSHD2 cells. We did not observe any overlap between enhancers deprived of H3K27me3 with enhancer harboring a differential DNA methylation profile in fibroblasts. Overall, changes in H3K27me3 distribution in SMCHD1-deficient muscle cells uncovers opposite functions for this protein: in condensation of constitutive heterochromatin while protecting poised and repressed chromatin against deposition of the repressive mark.

### SMCHD1 regulates H3K27me3 and CTCF distribution at HOX genes clusters

We first looked more closely at loci regulated by Smchd1 in the mouse, starting with the *HOX* clusters (Jansz et al., 2018). In BAMS or FSHD2 muscle cells, several *HOX* genes are differentially expressed compared to controls (FDR<0.05) with up-regulation of proximal *HOX* in BAMS and FSHD2 cells but down-regulation of distal genes (A10, C10 and D10) in FSHD2 (Figure S11C). H3K27me3 is highly enriched at all *HOX* loci in controls with a marked peak demarcation at loci borders. In BAMS and FSHD2 cells, large regions are depleted of this mark with no massive difference between cells carrying a gain (BAMS) or loss (FSHD2) of ATPase activity (Figure 5C, D; Figure S11A, B). In the mouse, absence of Smchd1 has been associated with a gain in CTCF (Jansz et al., 2018). We did not evidence a massive gain in CTCF binding across *HOX* clusters in BAMS and FSHD2 cells, except in the vicinity of *HOXA10* and *HOXC10* genes.

Overall, our work uncovers some overlapping functions for mouse and Human SMCHD1 in the distribution of H3K27me3 at large clusters encoding master developmental genes. However, SMCHD1 deficiency is not necessarily associated with a gain in CTCF binding, at least in cells where mutation affects the ATPase activity of the protein.

### SMCHD1 regulates biological pathways related to BAMS or FSHD2 phenotype

BPs associated with H3K4me3 peaks highlight the implication of SMCHD1 in non-coding RNAs regulation, chromatin organization and development with some overlaps between H3K4me3 and H3K27me3. BPs overrepresented for CTCF and H3K27me3 peaks are highly relevant to the diseases phenotype, *i*.*e*, related to cell migration in BAMS (Figure 6A,B) and muscle function in FSHD2 (Figure 6A,C). By comparing the list of genes showing a differential H3K27me3 profile to the list of genes differentially expressed in FSHD2 muscle fibers (Laberthonniere et al. Submitted), we retrieved 30 genes (table S11) with H3K27me3 peaks mapping to active or poised promoters, repressed chromatin or enhancers. Interestingly, all share the GVGGMGG motif (pvalue 9.42e^-6^), a binding site for the ETF (TEAD2) transcription factor, that binds tissue-specific enhancers and implicated in the regulation of neural crest cells, myogenic precursors (Kaneko et al., 2007) and muscle regeneration (Zhao et al., 2006). We then compared the list of genes near CTCF peaks (Figure 6D,E) to the list of genes differentially expressed in FSHD2 and BAMS muscle cells (Laberthonniere et al. Submitted). We retrieved 167 genes in FSHD2 (Figure 6D) and 665 in BAMS (Figure 6E). GO terms related to the corresponding BPs appear are also consistent with the respective disease phenotypes, i.e. sarcomeric structure in FSHD2, cartilage and sensory organ development in BAMS. Altogether, these data demonstrate that SMCHD1 acts on the conformation of chromatin of genes involved in lineage specification and relevant to the typical phenotypes associated with either FSHD2 or BAMS.

**Figure 6.**
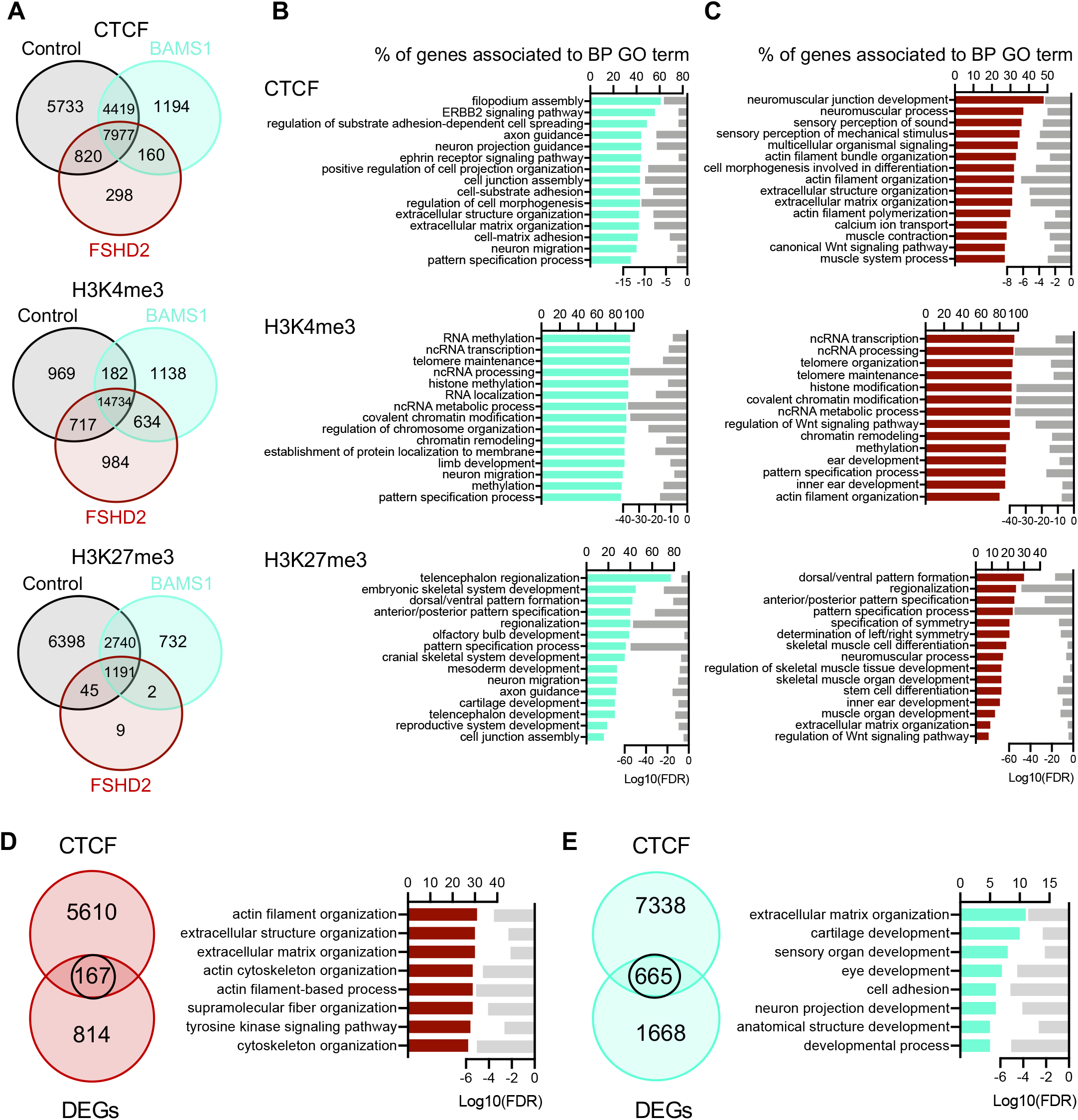
Differential chromatin profiling of HOX genes and distribution of CTCF peaks in SMCHD1-deficient muscle cells. **A**. Venn diagrams for comparison of peaks enriched in Controls, BAMS and FSHD2 hiPSC-derived muscles for CTCF, H3K4me3 or H3K27me3. **B**. GO tems for Biological pathways (BP) corresponding to peaks enriched in CTCF, H3K4me3 or H3K27me3 enrichment in BAMS (cyan) cells. Cyan bars correspond to the number of genes corresponding to the different BP, Light grey bars in the right represent the enrichment score (Log10 of False Discovery Rate) for each GO-term. **C**. GO terms for BP corresponding to peaks enriched in CTCF, H3K4me3 or H3K27me3 enrichment in FSHD2 cells. Red bars correspond to the number of genes corresponding to the different BP, Light grey bars in the right represent the enrichment score (Log10 of False Discovery Rate) for each GO-term. **D**. Venn diagrams for comparison of genes that are differentially expressed in FSHD2 muscle fibers derived from hiPSCs compared to controls with a fold-change >2 and <-2 and an FDR <0.05 and CTCF peaks. Biological pathways corresponding to the 167 overlapping genes were analyzed using g.profiler. Bar plots in the left represents the percentage of genes that are differentially expressed and associated with a GO-term shown in the right column. Light grey bars in the right represent the enrichment score (Log10 of False Discovery Rate) for each BP. **E**. Venn diagrams for comparison of genes that are differentially expressed in BAMS muscle fibers derived from hiPSCs compared to controls with a fold-change >2 and <-2 and an FDR <0.05 and CTCF peaks. Biological pathways corresponding to the 665 overlapping genes were analyzed as described above.

### CTCF binding is increased in the total absence of SMCHD1

We have shown that mutations in SMCHD1 ATPase domains affect distribution of H3K27 trimethylation but are not accompanied by an increase in CTCF binding as in KO mice (Jansz et al., 2018). To further investigate the difference between SMCHD1 deficiency and a total absence of the protein on CTCF distribution in human cells, we analyzed publicly available ChIP-Seq data (GEO, GSM1130654) where SMCHD1 binding was determined by comparing HCT116 cells and HCT116 in which *SMCHD1* has been invalidated using a Zn finger nuclease (HCT116 KO). After filtering for repetitive DNA sequences, 526 SMCHD1 binding sites were found. Most of them are located at a distance of 50-500 kb from TSS consistent with an indirect long-distance effect on gene regulation. By artificial extension of SMCHD1 peaks coordinates (± 10 kb from the peak), we obtained a list of genes in the vicinity of these peaks and compared this list to fibroblasts DEGs. We selected 24 DEGs with 14/24 harboring a CTCF site in their vicinity. We further characterized three of them with insulator features (*BET1L, NCAM2* and *SEMA5A*) together with one gene close to a putative SMCHD1 peak that does not overlap with CTCF binding (*WASH7PL*; no associated chromatin feature). By ChIP-qPCR, we confirmed a slight SMCHD1 enrichment at *SEMA5A* but did not detect any significant binding at BET1L, NCAM2 and WASH7PL (Figure S12). However, we noticed that SMCHD1 knock-out significantly increases CTCF occupancy (pvalue < 0.00001) at all four sites indicating that SMCHD1 might interfere, at least indirectly with the binding of the CTCF insulator protein. However, given the absence of CTCF enrichment in cells carrying a mutation in the ATPase domain, we concluded on a different impact of SMCHD1 deficiency or absence on CTCF distribution and chromatin architecture.

### SMCHD1 regulates gene expression *in cis*

We have evidenced a role for SMCHD1 in the regulation of enhancers episignatures and in shielding against the deposition of chromatin repressive marks. We then proceeded to functional testing a number of SMCHD1 target sequences in gene regulation and insulation using transient and stable reporter gene expression assays. Transient reporter assays will evaluate enhancer or promoter activity whereas stable insertions will reveal position effects. We first started with the D4Z4 macrosatellite and tested subregions bound by SMCHD1 (Dion et al., 2019), that display differential methylation profile in patients (Gaillard et al., 2014; Roche et al., 2019) (figure S13A) on transient Luciferase expression. In a vector lacking the SV40 enhancer element, the proximal part of D4Z4 differentially methylated in *SMCHD1*-mutated cells (DR1 and DR1 subfragments, up to CG 21) activates luciferase expression in an SMCHD1-dependent manner (Figure 7A,B) but has no activity in a promoter-less reporter (Figure S13B). We did not evidence any enhancer activity for the different HOX DMR that bind SMCHD1 in HEK cells (Figure 7C). However, we observed an increased luciferase expression in the absence of SMCHD1 for HOXB2, B6 and HOXC4 DMRs showing that SMCHD1 regulates gene expression by binding to these DMRs (Figure S8). The same observation was made for the potential binding site near *SEMA5A* identified as a potential SMCHD1 binding site (Figure 7D) but not for *NCAM2, BET1L*. We concluded that in transient assays, SMCHD1 contributes either to transcription activation or silencing depending on its target sequence.

**Figure 7.**
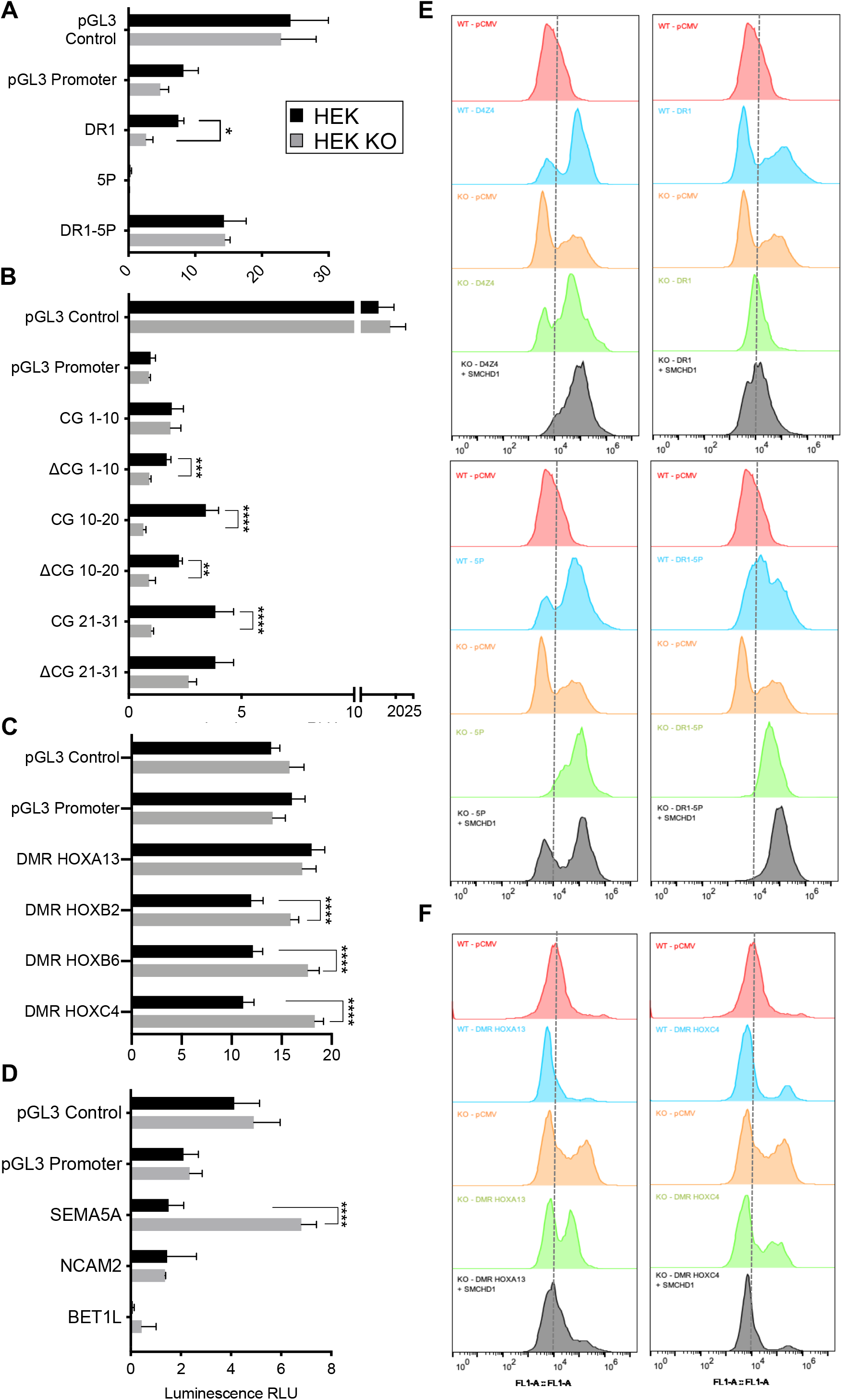
Depending on the genomic context, SMCHD1 contributes to gene silencing or protects against position effects. **A-D**. For the different pGL3 constructs fragments corresponding to the regions that are differentially methylated (**A**-**C**) or corresponding to SMCHD1 binding sites (**D**) were cloned downstream of the Luciferase reporter gene in vectors lacking an enhancer (pGL3 promoter). Firefly luciferase expression was determined 48 hrs post-transfection of the different constructs in HEK 293 and HEK *SMCHD1* KO cells. The pGL3 control vector was used as transfection control. Firely luciferase levels were normalized to expression of the Renilla luciferase used as transfection control. Values corresponding to the normalized luciferase activity (expressed in relative luminescence units, RLU) are the average of three independent assays, each realized as technical triplicates (n=9). Error bars represent standard error. Statistical significance was determined using a Mann Whitney test, **** p value < 0.00001, *** p value <0.0001, ** p value< 0.001, * p value = 0.01). **A**. For the D4Z4 macrosatellite, regions that are differentially methylated in patients carrying a mutation in *SMCHD1* were tested (DR1, 5P. A scheme of the D4Z4 repeat is presented in supplementary figure 13). **B**. The DR1 sequence contains 31 CpG sites. Different fragments corresponding to CG1-10, CG10-20, CG21-31) or deleted of 10 of these CG (ΔCG1-10, ΔCG10-20, ΔCG21-31) were tested. **C**. H*OX* genes DMR. **D**. Putative SMCHD1 binding sites overlapping or not with CTCF binding sites. **E-F**. For evaluation of protection against position effect, we used a vector carrying a Hygromycin resistance gene fused to the herpes simplex virus type 1 thymidine kinase suicide gene (*HyTK)* and an *eGFP* reporter gene, each driven by a CMV promoter (pr) (pCMV construct). Sequences to be tested are cloned downstream of the *eGFP* gene. Constructs were linearized and transfected into HEK or HEK KO Cells. Stable eGFP expression was measured by flow cytometry (FACS) for an extended period of time in cells grown in the presence of Hygromycin B. Representative spectra of the % of eGFP positive cells a presented. eGFP expression level is the average of 5 measurements from day 18 to day 40 post-transfection, when *eGFP* expression reaches a plateau, for three independent assays ± S.D. For each condition, eGFP expression was compared to values obtained in cells transfected with the empty vector (pCMV). In HEK *SMCHD1* KO cells, eGFP expression was also measured 72 hrs after transfection of a SMCHD1 expression vector (grey curves) **E**. D4Z4 (left upper panel), DR1-5P (right upper panel), DR1 (left lower panel) and 5P (right lower panel) were cloned downstream of the eGFP reporter. **F**. Results obtained for the HOXA13 (left) and HOXC4/5/6 (right) DMRs.

### SMCHD1 is able to protect against position effect variegation in human tissues

We then tested the impact of SMCHD1 on Position Effect Variegation (PEV) after stable transfection of our previously described reporter system, using the D4Z4 insulator as a reference (Ottaviani et al., 2009a; Ottaviani et al., 2010) (Figure 7E). As in C33A cells, D4Z4 protects against PEV in HEK293 cells as indicated by the increased proportion of eGFP-positive cells (Figure 7E, blue shading). This D4Z4-dependent protection against PEV is strongly increased when SMCHD1 is overexpressed in HEK KO cells (Figure 7E, grey shading) showing that SMCHD1 contributes to D4Z4 anti-PEV activity. DR1 harbors a moderate anti-PEV effect that is not improved by *SMCHD1* overexpression. The 5P region or the DR1-5P combination increases the percentage of cells expressing the eGFP reporter revealing a protection against position effect that is dependent on SMCHD1.

As in luciferase assays, HOXA13 DMR has no effect on the eGFP reporter gene expression (Figure 7F). By increasing the percentage of eGFP-positive cells, HOXB2 DMR acts as an activating element partially regulated by SMCHD1 (KO cells, orange shading, Figure S13D). For HOXB6 (Figure S13D) and HOXC4/C5/C6 (Figure 7F), we observed a slight increase in the percentage of eGFP-positive cells, augmented in the absence of SMCHD1 indicating that SMCHD1 negatively regulates HOXB6 and HOXC4/C5/C6 DMRs as in transient assays consolidating previous findings. Regarding SMCHD1 binding sites identified in HCT116/HCT116 KO cells, similar profiles were obtained for NCAM2, BET1L and WASH7PL with a marked increase in eGFP-positive cells in HEK KO cells that is counter balanced upon *SMCHD1* re-expression (Figure S13D). These different regions repress gene expression in a SMCHD1-dependent manner while SEMA5A effect on gene expression is not mediated by SMCHD1. Overall, these results highlight multiple roles for SMCHD1 as a *cis-*activator (D4Z4-DR1, *SEMA5A*), or as a moderate (*HOXB2, NCAM2, WASH7PL*) to strong (*HOXA13, HOXB6, HOXC4/C5/C6, BET1L*) repressor of gene expression.

## Discussion

First identified as a regulator of variegation in the mouse (Blewitt et al., 2005), Smchd1 is required for proper X inactivation and regulates large gene clusters such as *Hox* or *Pcdh* loci and imprinted genes chromatin (Blewitt et al., 2008; Brideau et al., 2015; Gdula et al., 2019; Gendrel et al., 2012; Jansz et al., 2018; Mason et al., 2017; Massah et al., 2014; Wang et al., 2018; Wanigasuriya et al., 2020). At the 2 cell stage, Smchd1 shields against active Tet-dependent DNA demethylation (Huang et al., 2021) and plays a predominant role in autosomal imprinting through a maternal effect (Ruebel et al., 2019; Wanigasuriya et al., 2020). However, the role of this protein in human tissues remain partially understood despite its implication in a number of distinct rare genetic diseases linked to heterozygous *SMCHD1* mutation (Gordon et al., 2017; Kinjo et al., 2020; Lemmers et al., 2012; Shaw et al., 2017).

To define SMCHD1 function in the epigenetic regulation of the human genome and its implication in FSHD and BAMS phenotype, we performed genome-wide DNA methylation and chromatin profiling in cells from patients carrying a heterozygous mutation in this gene. Compared to the fully penetrant effect of gene invalidation in the mouse, *SMCHD1* mutation in BAMS and FSHD2 acts on protein dosage or function with gain of function mutations in the majority BAMS patients and haploinsufficiency or loss of function in FSHD2. A key difference between mouse and Human also resides in the absence of BAMS and FSHD2-like phenotype in the mouse and the lack of a clear evidence for X inactivation defect in BAMS and FSHD2 patients (Dion et al., 2019) as both sexes are equally affected. By focusing on the role of this non-canonical Cohesin in the regulation of autosomes, we showed that SMCHD1 shapes chromatin at different levels depending on the genomic context. As in the mouse, SMCHD1 contributes to *de novo* methylation and is dispensable for methylation maintenance but its occupancy is not correlated to the level of DNA methylation (Dion et al., 2019; Gdula et al., 2019; Huang et al., 2021). We further showed that SMCHD1 is required for the maintenance of heterochromatin but also shields against spreading of repressive chromatin marks suggesting a role in chromatin insulation. Furthermore, SMCHD1 regulates the episignature of a number of enhancers orchestrating expression of genes related to development and cell differentiation circuits.

Mouse Smchd1-deprived cells harbor increased binding of the CTCF architectural protein (Gdula et al., 2019; Jansz et al., 2018; Wang et al., 2018). Moreover, absence of Smchd1 trigger long-range extra TADs interactions (Jansz et al., 2018) and changes in the Xi topology (Gdula et al., 2019; Jansz et al., 2018; Wang et al., 2018). In BAMS and FSHD2 patient’s muscle cells, we did not evidence any increase in CTCF binding but an overall decrease in CTCF peaks. Based on our previous results showing an increased binding of SMCHD1 to D4Z4 of 2-fold in BAMS and up to 6-8-fold in FSHD2 patient’s cells (Dion et al., 2019), we speculate that some of the mutations, especially those affecting the ATPase domain might increase the affinity of the homo-dimeric SMCHD1 complex to chromatin or hamper its removal, subsequently restraining CTCF access. Alternatively, the titration of functional SMCHD1 complexes might also lead to a variable relative occupancy between SMCHD1 and CTCF from cell to cell. In human cells, SMCHD1 contributes to chromatin insulation by acting as a boundary against repressive chromatin, in particular by regulating the chromatin boundary activity of D4Z4 (Ottaviani et al., 2009a; Ottaviani et al., 2010).

We also uncovered a role for SMCHD1 in the regulation of enhancers at two levels, H3K27me3 deposition and DNA methylation. Multiple roles are thus attributable to SMCHD1 that acts either as a *cis-*activator (D4Z4-DR1, *SEMA5A*), or as a moderate (*HOXB2, NCAM2, WASH7PL*) to strong (*HOXA13, HOXB6, HOXC4/C5/C6, BET1L*) repressor of gene expression. As Smchd1 protects against Tet-dependent demethylation (Huang et al., 2021), it remains to be determined whether the increased proportion of hypomethylated enhancers in BAMS dans FSHD2 cells is due to SMCHD1 inability to counteract active demethylation at enhancers related to during cell fate transition and cell lineage specification.

As autosomal dominant SMCHD1 mutations impact stem cells properties as well as post natal health(Schall et al., 2019), the diverse phenotypes of *SMCHD1*-mutated patients and animal models might be the consequence of pleiotropic effects triggered by changes in chromatin boundaries and variegation in gene expression of loci involved in development and differentiation. Based on our findings, one can speculate that changes in the epigenetic signature of a number of predicted SMCHD1-dependent enhancers might induce promiscuous or precocious activation/repression of genes located at a long-distance from these regulatory elements with variable impacts depending on the type of mutation. To date, the molecular mechanisms specifying which of the thousands of CTCF sites are regulated by DNA methylation and involved in cell differentiation remain elusive. This emphasizes the need of understanding how SMCHD1 regulates DNA methylation and deposition of repressive chromatin marks and subsequently, overlap between SMCHD1 and CTCF in guiding genome topology. Very interestingly, CTCF has been hypothesized to have co-evolved with *Hox* genes as a regulator of spatio-temporal gene expression (Heger et al., 2012). Furthermore, hERV have spread CTCF binding sites in the human genome, an interesting observation considering the role of SMCHD1 in regulating the D4Z4 macrosatellite repeat encoding the DUX4 transcription factor required for zygotic genome activation and regulation of hERVs (De Iaco et al., 2017; Hendrickson et al., 2017).

Together with previous reports, our work confirms a key role for SMCHD1 in the functional partitioning of the mammalian genome with consequences in diseases. Boundary disruption has been linked to cancer, diseases of limb development or linked to triplet expansions (Flavahan et al., 2016; Hnisz et al., 2016; Lupianez et al., 2015; Sun et al., 2018). We expand here the list of diseases associated with topological changes by showing that mutations in *SMCHD1* disrupt a subset of chromatin boundaries and contribute to the mechanisms restricting promiscuous enhancers interactions. Reminiscent of the role of Dxz4 in the mouse (Gdula et al., 2019; Wang et al., 2018), this work also opens new avenues on the role of repetitive elements in the formation of topological domains and chromatin boundaries involving repetitive DNA elements such as D4Z4 (Francastel and Magdinier, 2019; Ottaviani et al., 2010; Ottaviani et al., 2009b; Robin et al., 2015),

In conclusion, in patients carrying a heterozygous *SMCHD1* mutation, SMCHD1 deficiency is phenotypically less drastic that in KO mice. We thus hypothesize that titration of functional SMCHD1 dimers alters DNA and K27 methylation and long-range chromatin organization at a number of loci. This in turn triggers variegation in the expression of genes required for development, cell fate transition and tissue differentiation in rare genetic diseases linked to heterozygous *SMCHD1* mutations. With a strong translational perspective but also on fundamental knowledge, our work further emphasizes the need of in-depth characterization of SMCHD1 functional domains that will enable us to decipher the respective consequences of germline mutations in diseases for genetic diagnostics and therapeutic interventions.

## Acknowledgements and Fundings

We are indebted and thank all patients for participating in this study. We acknowledge the GBiM genomics and bioinformatics core facility for RNA seq and the Marseille Stem Cells Core facility (MaSC) for stem cells. This study was funded by “Association Française contre les Myopathies” (AFM; TRIM-RD) and Fondation Maladies Rares. CD and CL were the recipient of a fellowship from the French Ministry of Education and FSH Society. MD is the recipient of a fellowship from the French Ministry of Education. The project leading to this publication has received funding from the Excellence Initiative of Aix-Marseille University-A*Midex, a French “investissement d’avenir programme” AMX-19-IET-007 through the Marseille Maladies Rares (MarMaRa) Institute.

