## Supplementary figures and text for "SMCHD1 variants may induce variegated expression in Facio Scapulo Humeral Dystophy and Bosma Arhinia and microphtalmia syndrome"

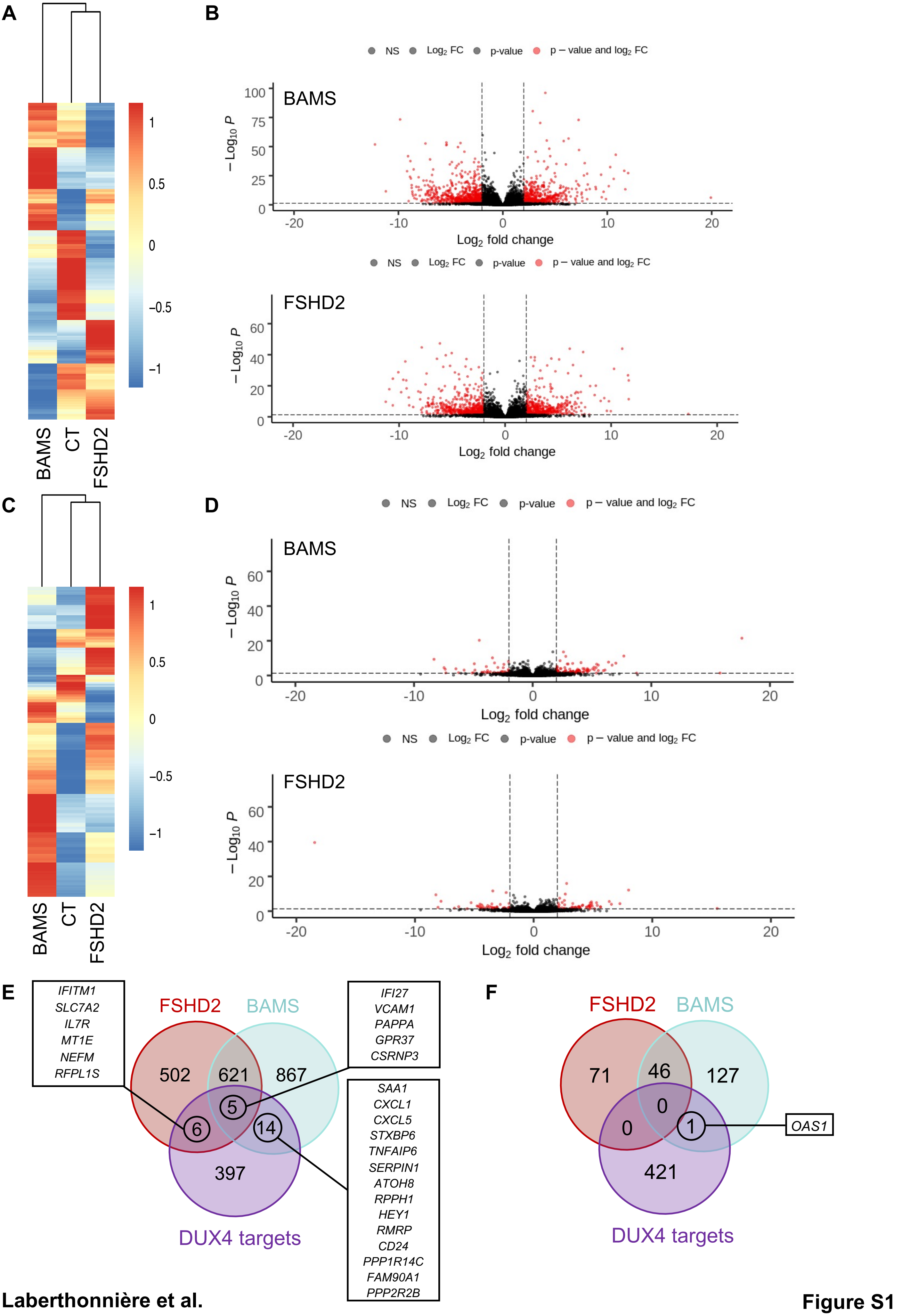


**Laberthonnière et al.**

**Figure S1**

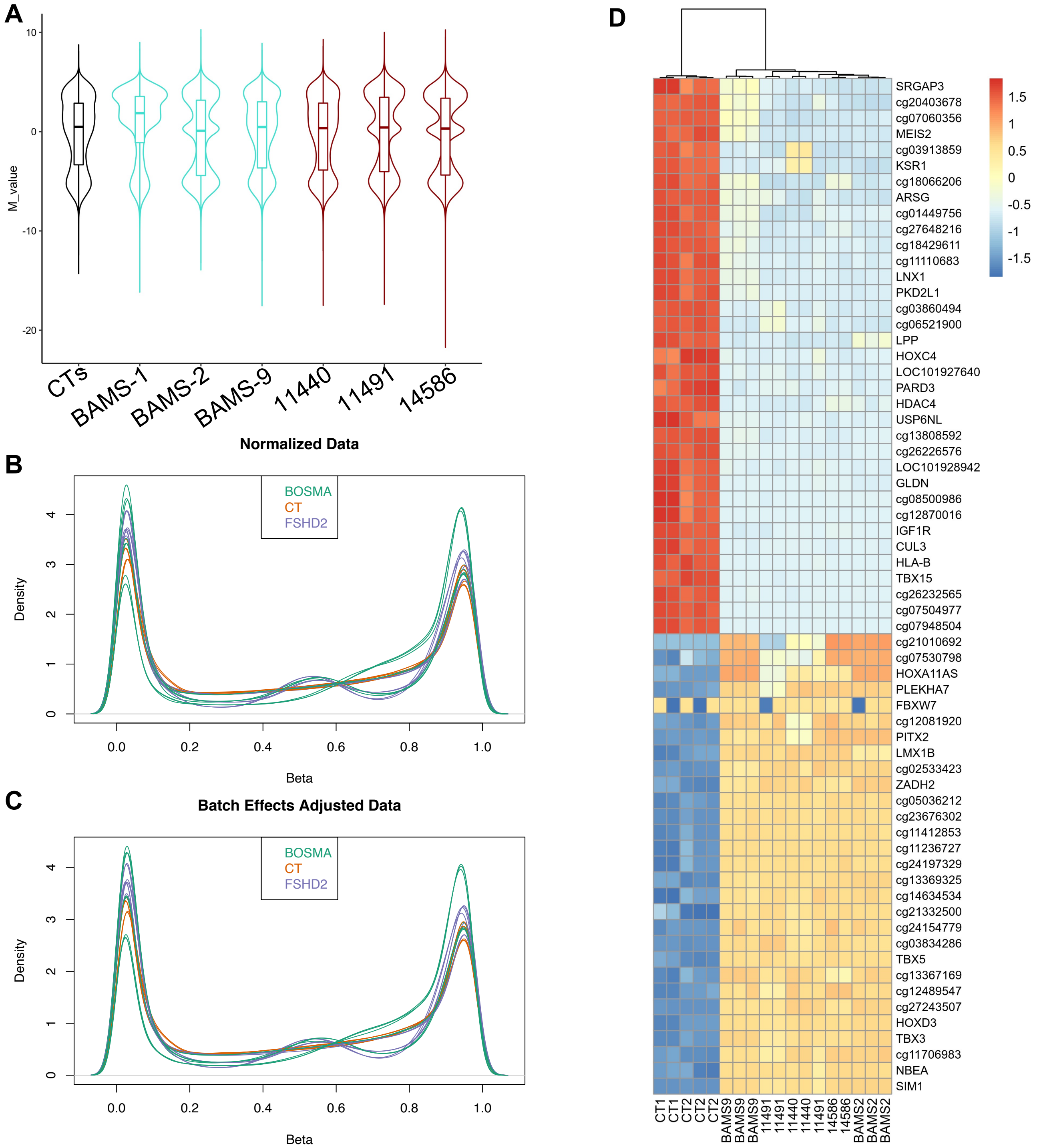


**Laberthonnière et al.**

**Figure S2**

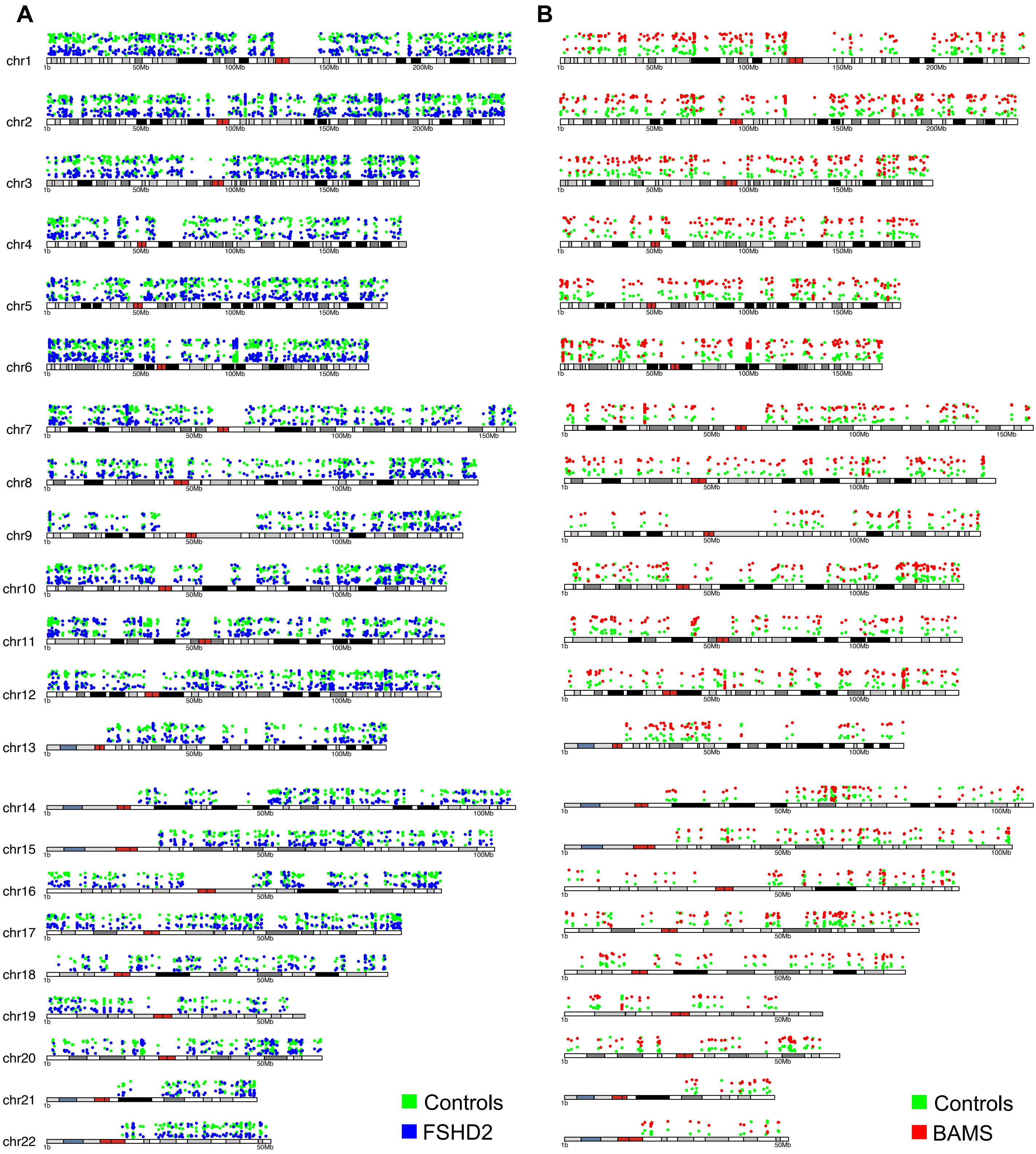
A
B
chr1
chr2
chr3
chr4
chr5
chr6
chr7
chr8
chr9
chr10
chr11
chr12
chr13
chr14
chr15
chr16
chr17
chr18
chr19
chr20
chr21
chr22
Controls
FSHD2
Controls
BAMS

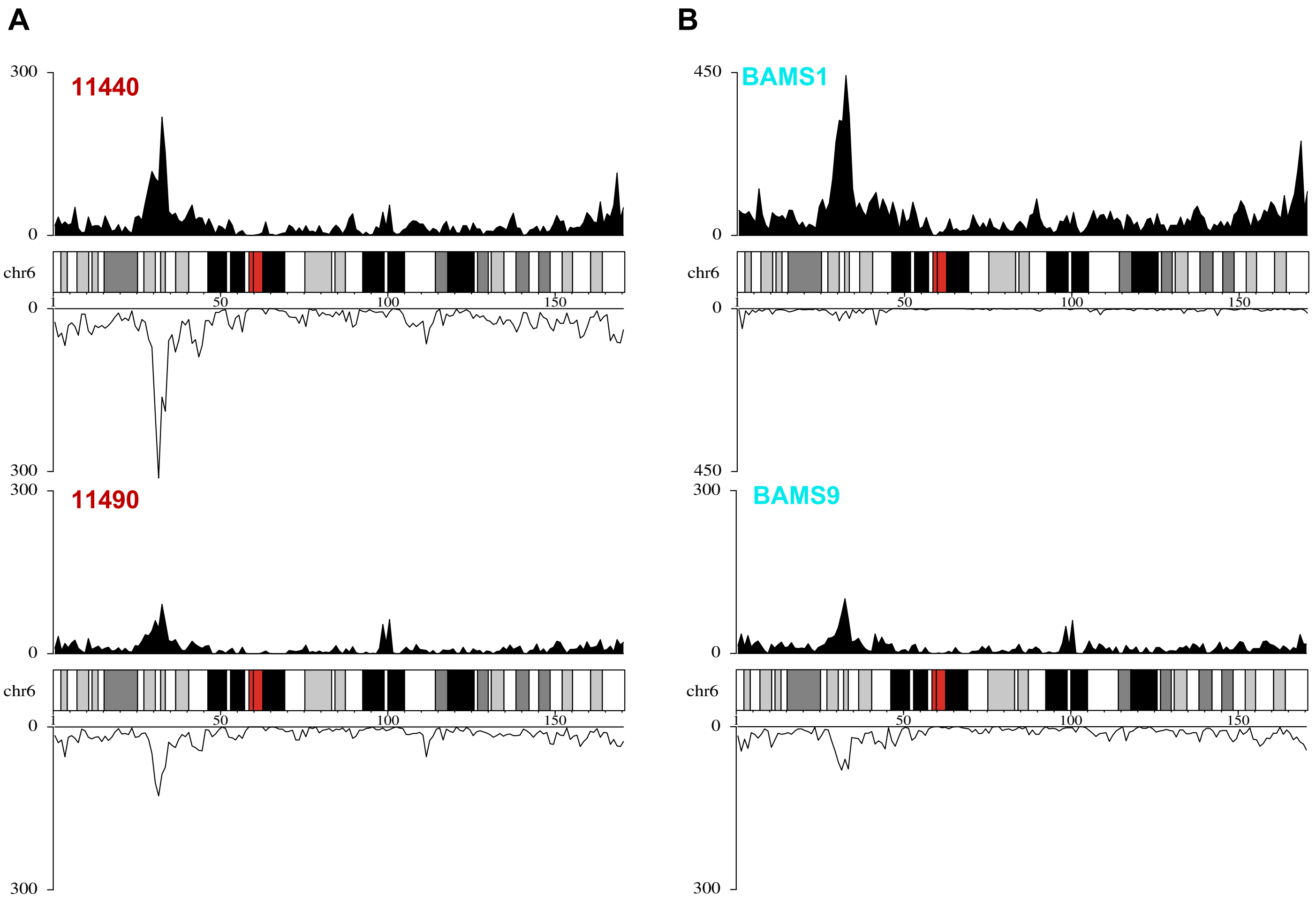


Laberthonnière et al.

Figure S4

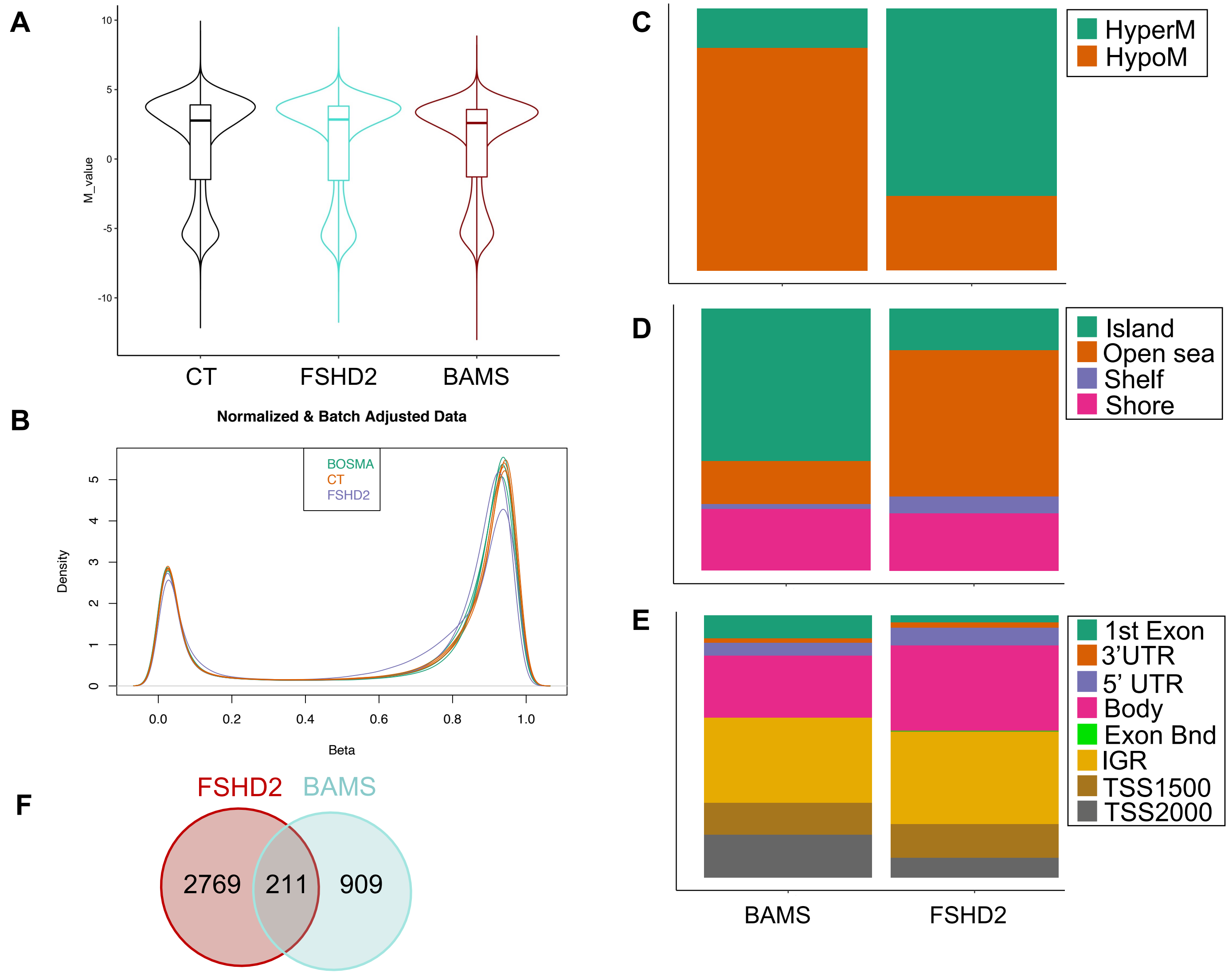


Laberthonnière et al.

Figure S5

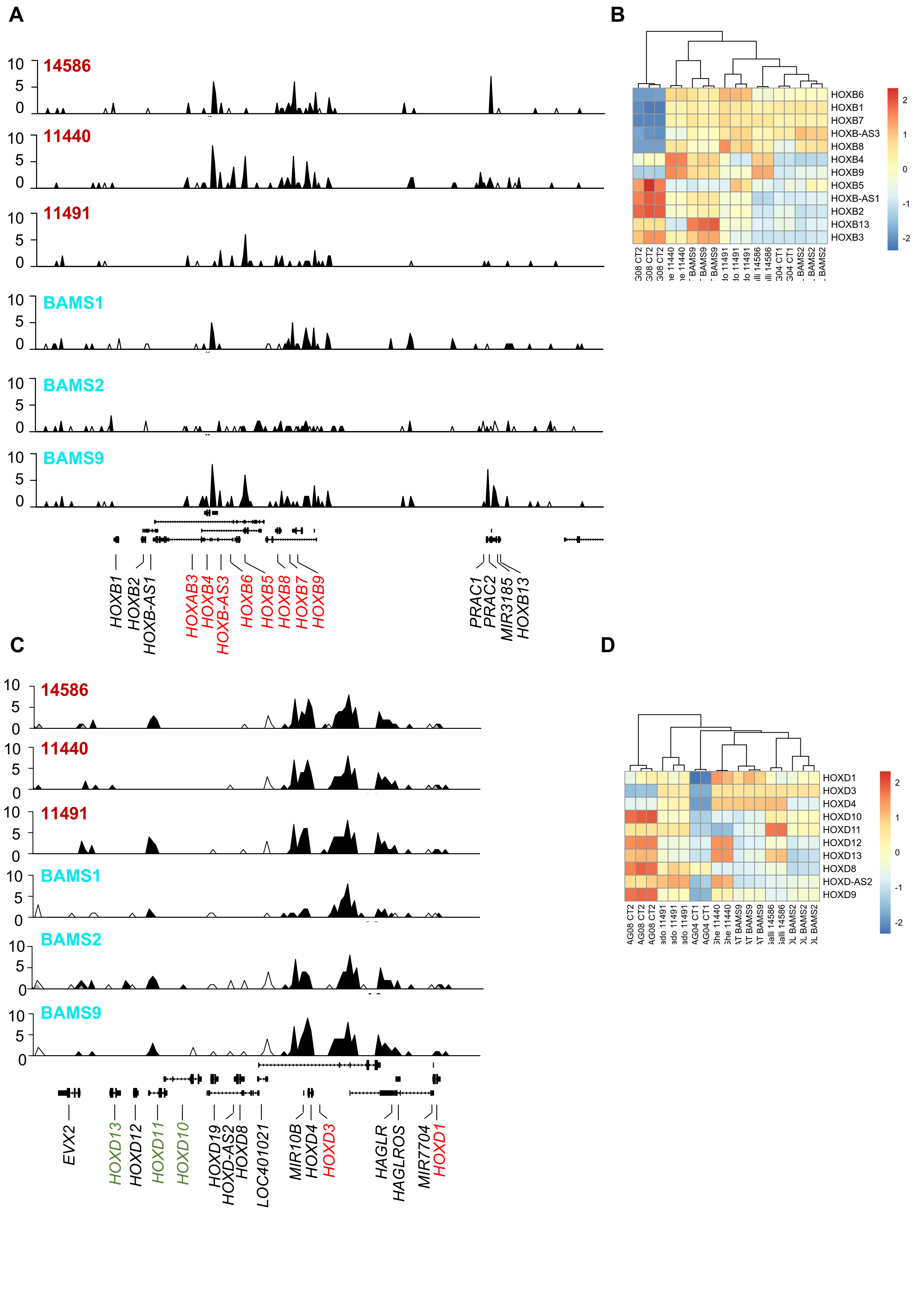
A
14586
11440
11491
BAMS1
BAMS2
BAMS9
HOXB1
HOXB2
HOXB-AS1
HOXAB3
HOXB4
HOXB-AS3
HOXB6
HOXB5
HOXB8
HOXB7
HOXB9
PRAC1
PRAC2
MIR3185
HOXB13
B
HOXB6
HOXB1
HOXB7
HOXB-AS3
HOXB8
HOXB4
HOXB9
HOXB5
HOXB-AS1
HOXB2
HOXB13
HOXB3
C
14586
11440
11491
BAMS1
BAMS2
BAMS9
EVX2
HOXD13
HOXD12
HOXD11
HOXD10
HOXD19
HOXD-AS2
HOXD8
LOC401021
MIR10B
HOXD4
HOXD3
HAGLR
HAGLROS
MIR7704
HOXD1
D
HOXD1
HOXD3
HOXD4
HOXD10
HOXD11
HOXD12
HOXD13
HOXD8
HOXD-AS2
HOXD9

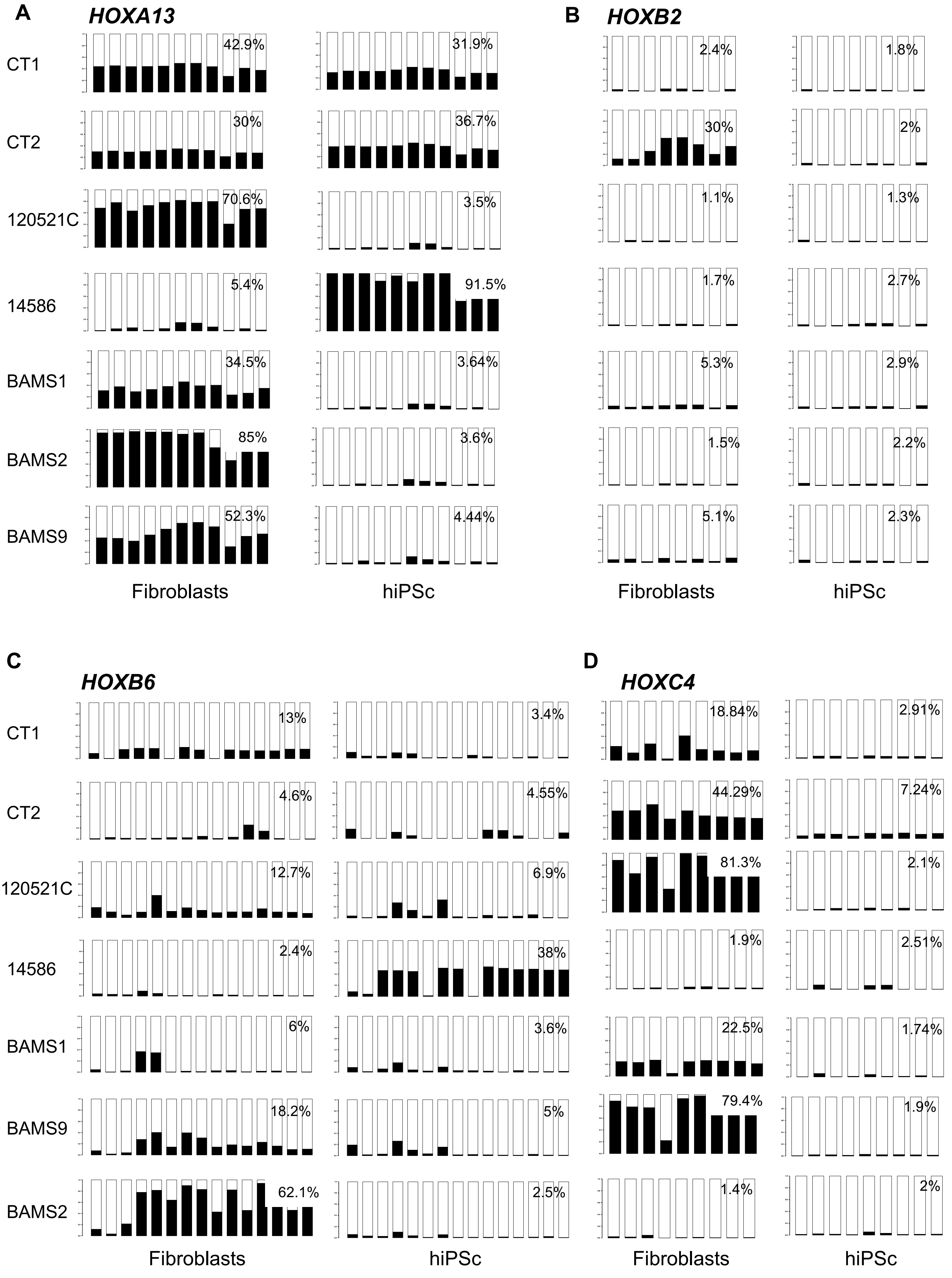


**Laberthonnière et al.**

**Figure S7**

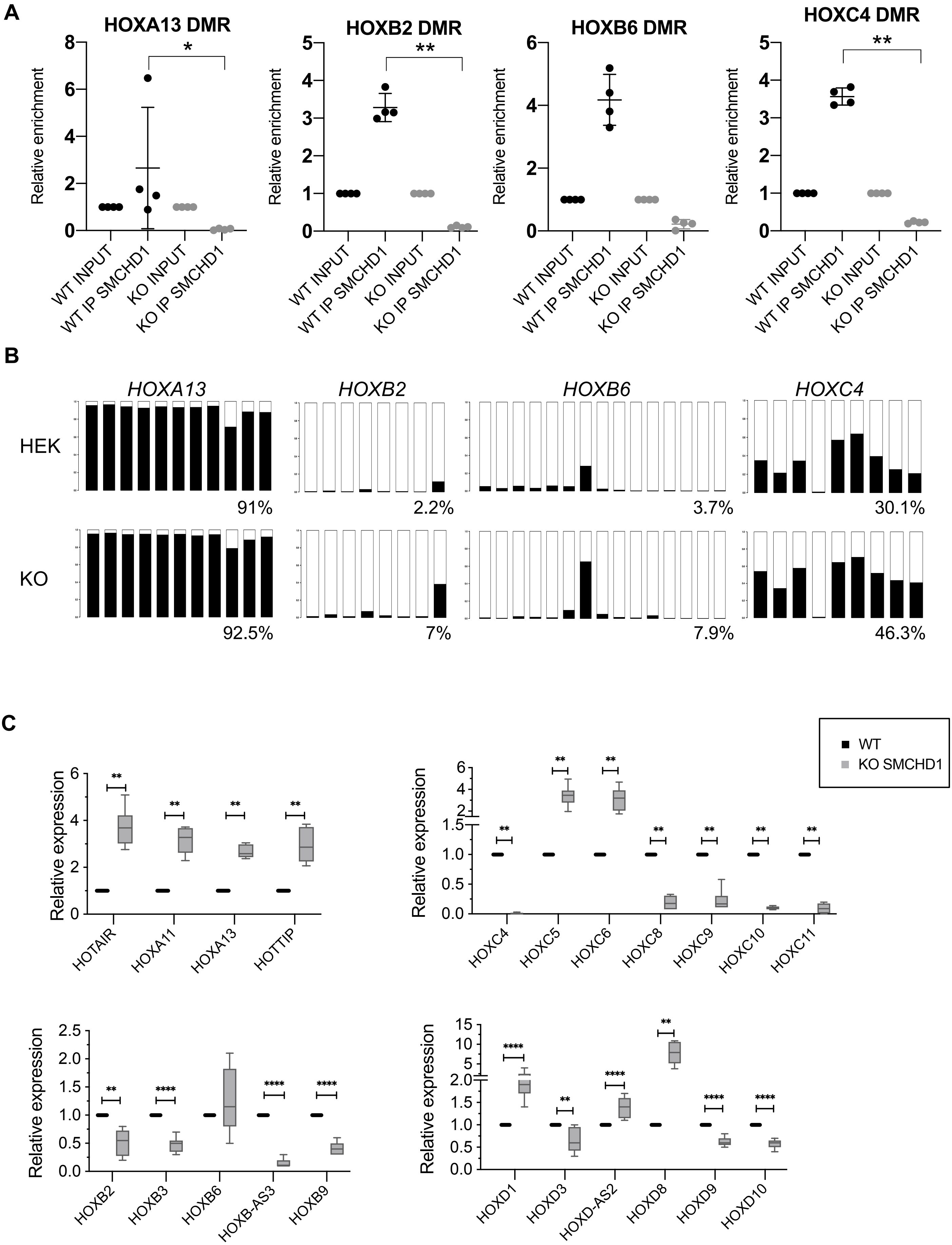


Laberthonnière et al.

Figure S8

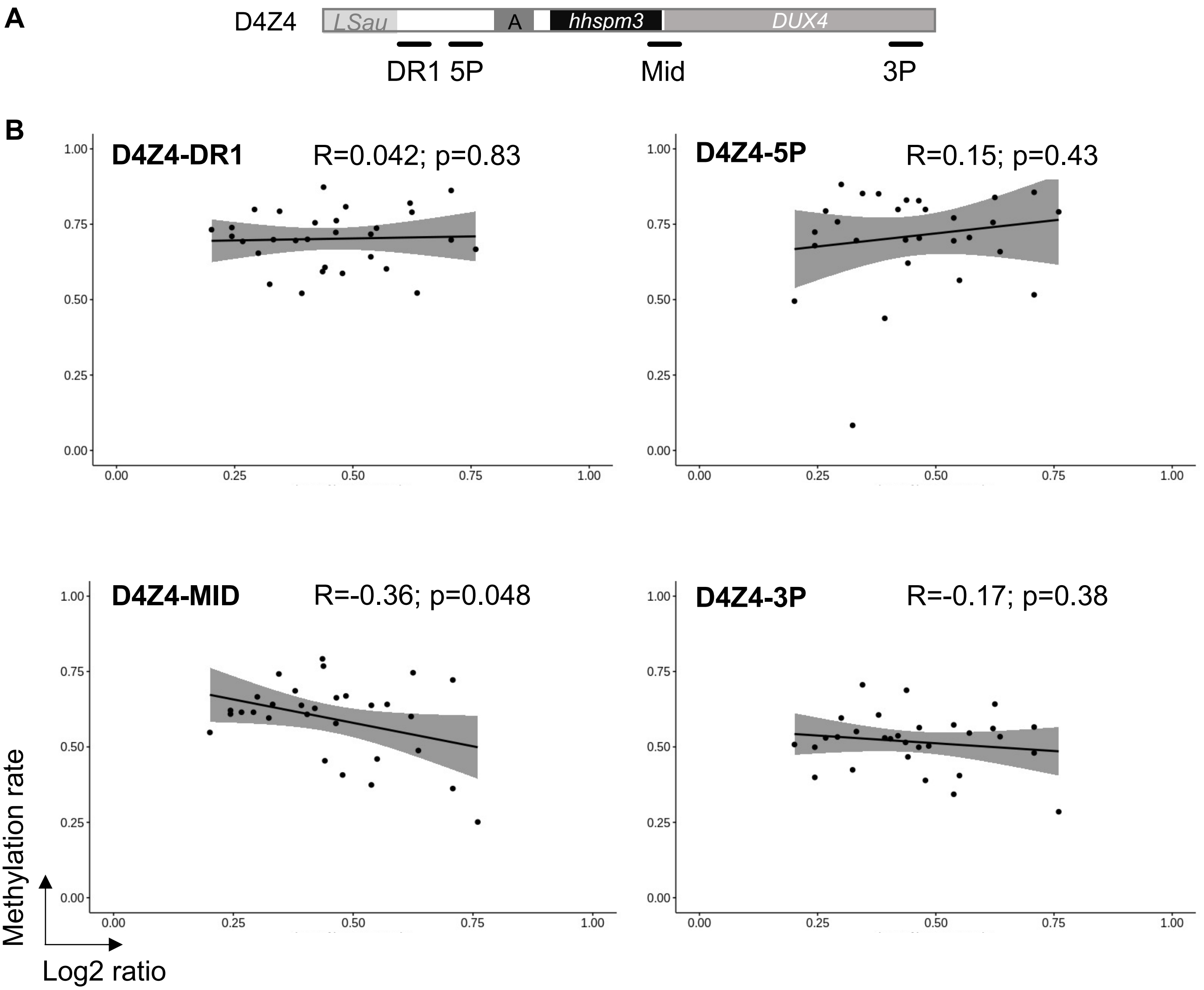
A
D4Z4
LSau
A
hhspm3
DUX4
DR1
5P
Mid
3P
B
D4Z4-DR1
R=0.042; p=0.83
D4Z4-5P
R=0.15; p=0.43
D4Z4-MID
R=-0.36; p=0.048
D4Z4-3P
R=-0.17; p=0.38
1.00
0.75
0.50
0.25
0.00
Methylation rate
Log2 ratio

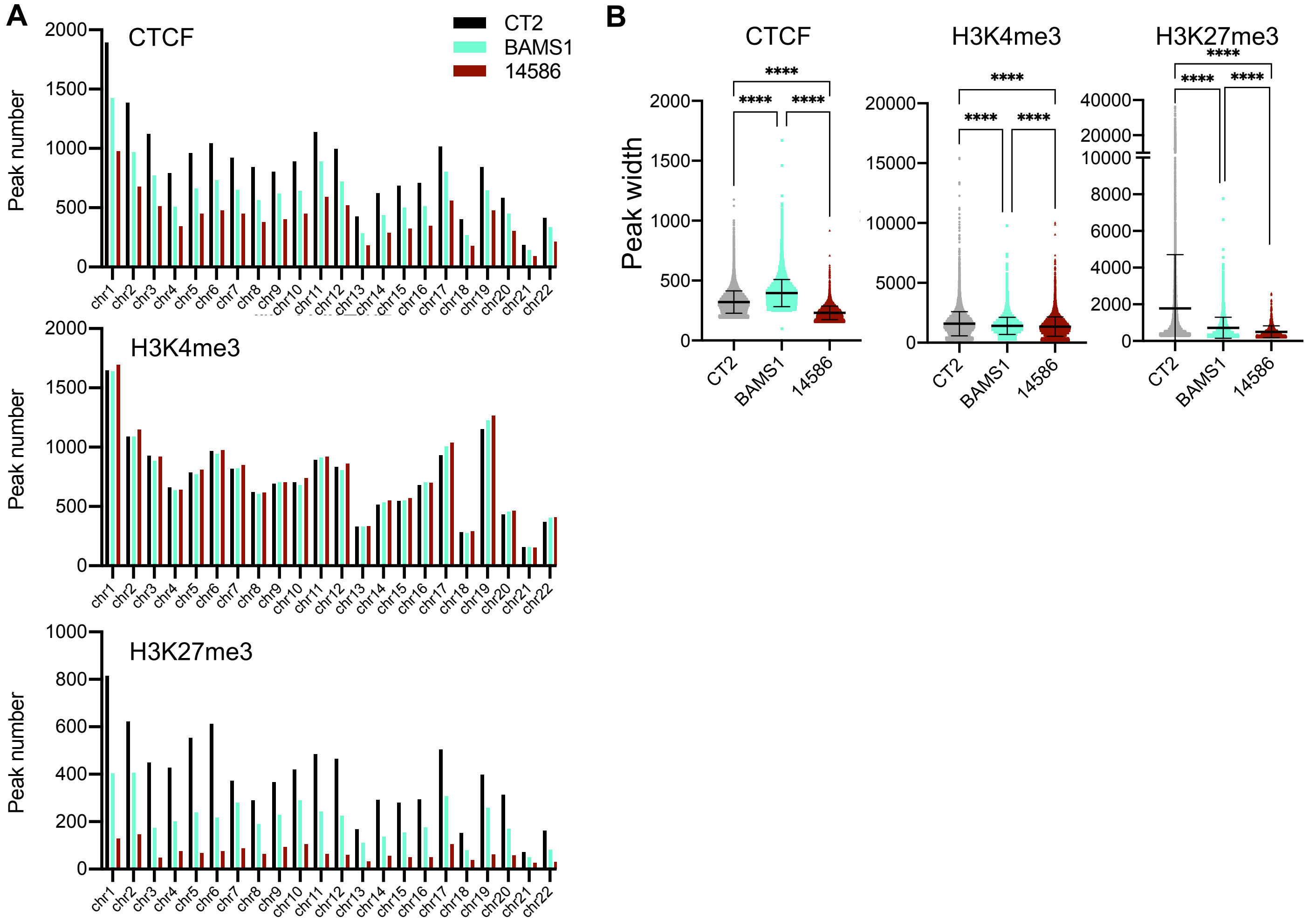


Laberthonnière et al.

Figure S10

A

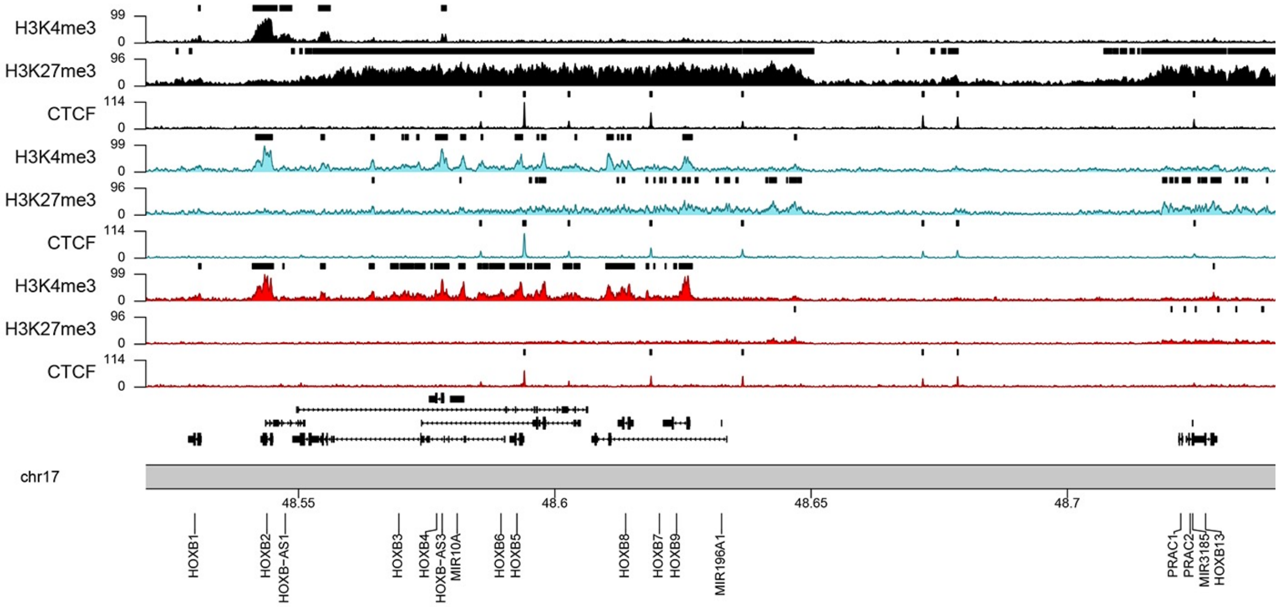


B

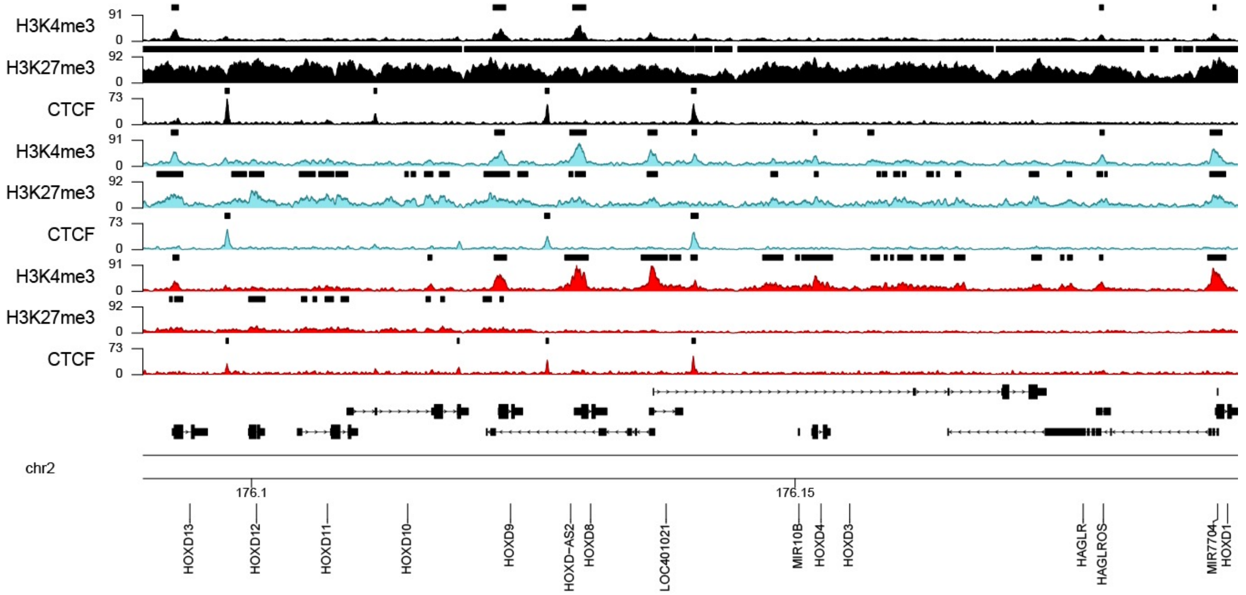


C

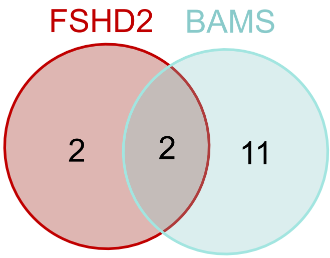


| | FC FSHD2 |
|---|---|
| HOXA-AS3 | -2.58 |
| HOXB9 | -2.29 |

| | FC FSHD2 | FC BAMS |
|---|---|---|
| *HOXB1* | -4.14 | -4.55 |
| *HOXD3* | -2.15 | -2.66 |

| | FC BAMS |
|---|---|
| HOXA-AS2 | -3.37 |
| HOXA10 | 2.56 |
| HOXA2 | -2.73 |
| HOXA3 | -2.09 |
| HOXA4 | -2.85 |
| HOXB2 | -2.04 |
| HOXB3 | -2.26 |
| HOXC10 | 4.26 |
| HOXC5 | -3.26 |
| HOXC8 | -2.90 |
| HOXD10 | 7.09 |

**Laberthonnière et al.** **Figure S11**

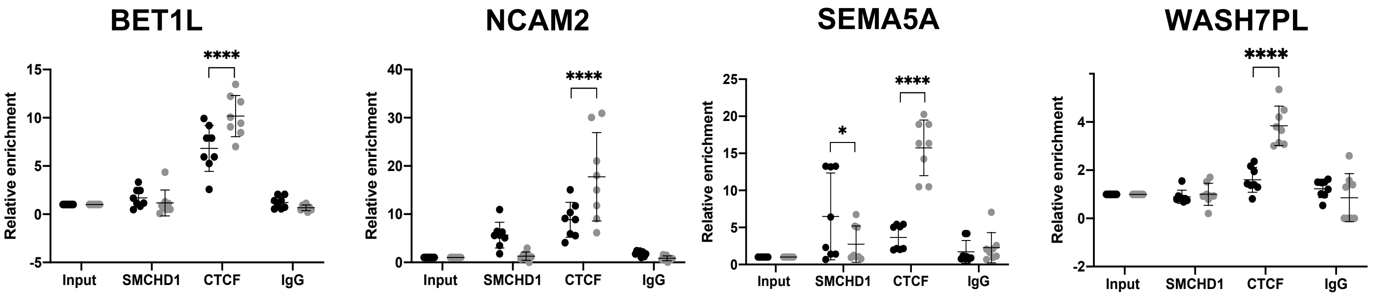


Laberthonnière et al.

Figure S12

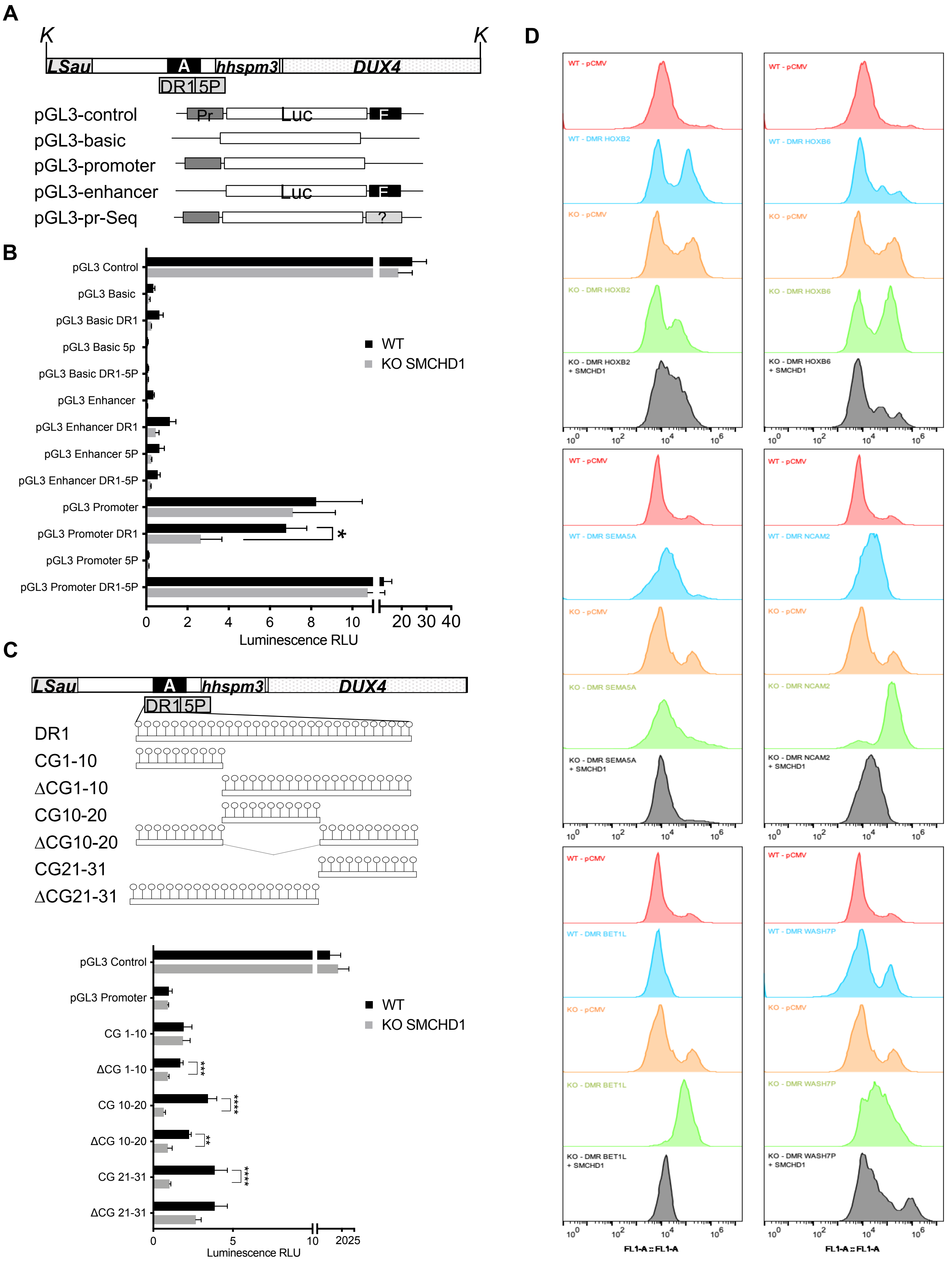
A
K
K
LSau
A
hhspm3
DUX4
DR1
5P
pGL3-control
Pr
Luc
E
pGL3-basic
pGL3-promoter
pGL3-enhancer
pGL3-pr-Seq
?
B
pGL3 Control
pGL3 Basic
pGL3 Basic DR1
pGL3 Basic 5p
pGL3 Basic DR1-5P
pGL3 Enhancer
pGL3 Enhancer DR1
pGL3 Enhancer 5P
pGL3 Enhancer DR1-5P
pGL3 Promoter
pGL3 Promoter DR1
pGL3 Promoter 5P
pGL3 Promoter DR1-5P
*
WT
KO SMCHD1
0 2 4 6 8 10 20 30 40
Luminescence RLU
C
LSau
A
hhspm3
DUX4
DR1
5P
DR1
CG1-10
ΔCG1-10
CG10-20
ΔCG10-20
CG21-31
ΔCG21-31
pGL3 Control
pGL3 Promoter
CG 1-10
ΔCG 1-10
CG 10-20
ΔCG 10-20
CG 21-31
ΔCG 21-31
***
****
**
****
WT
KO SMCHD1
0 5 10 2025
Luminescence RLU
D
WT - pCMV
WT - DMR HOXB2
KO - pCMV
KO - DMR HOXB2
KO - DMR HOXB2 + SMCHD1
WT - pCMV
WT - DMR HOXB6
KO - pCMV
KO - DMR HOXB6
KO - DMR HOXB6 + SMCHD1
WT - pCMV
WT - DMR SEMA5A
KO - pCMV
KO - DMR SEMA5A
KO - DMR SEMA5A + SMCHD1
WT - pCMV
WT - DMR NCAM2
KO - pCMV
KO - DMR NCAM2
KO - DMR NCAM2 + SMCHD1
WT - pCMV
WT - DMR BET1L
KO - pCMV
KO - DMR BET1L
KO - DMR BET1L + SMCHD1
WT - pCMV
WT - DMR WASH7P
KO - pCMV
KO - DMR WASH7P
KO - DMR WASH7P + SMCHD1
$10^0$ $10^2$ $10^4$ $10^6$
FL1-A :: FL1-A
FL1-A :: FL1-A

### Supplementary information

#### Materials and Methods

#### Samples.

All individuals have provided written informed consent for the collection of samples and subsequent analysis for research purposes. The study was done in accordance with the Declaration of Helsinki. Controls are randomly chosen individuals selected in the same age range and sex representation as patients. Controls are neither carrier of any genetic mutation nor affected by any constitutive pathology. Samples are listed in Table S1 and described in (Dion et al., 2019).

#### Cell Culture.

The human embryonic kidney 293 cell line (CRL-1573) was obtained from ATCC. In HEK293 cells, *SMCHD1* expression was invalidated by transfection of Zn finger nucleases. Cells were provided by J. Déjardin. Primary fibroblasts and HEK293 cells were grown in DMEM with L-alanyl-L-glutamine (GlutaMAX$^{TM}$I), D-glucose and sodium pyruvate (Life technologies). Media were supplemented in with 10% fetal bovine serum (FBS, Gibco), 1% penicillin/streptomycin (P/S). Cells were grown at 37°C, 5% $CO_2$, in a humidified atmosphere. Passages were performed with 1X Trypsin-EDTA (Gibco™) twice a week.

#### Human iPSCs clones.

All iPSCs clones were derived from primary fibroblasts (Table S1) after transfection of different vectors by electroporation (pCXCLE-hOCT3/4-shp53-F (Addgene ref 27077), pCXLE-hSK (Addgene, ref 27078, encoding *SOX2* and *KLF4*), pCXLE-hUL (Addgene, ref 27080, encoding *L-Myc* and *LIN28*). All cells were described in (Badja et al., 2014; Dion et al., 2019).

#### Neuromuscular hIPSc differentiation

The hIPSc were differentiated into functional muscles according to the protocol described in (Delourme et al., 2020; Mazaleyrat et al., 2020). In brief, at day 0, clumps of iPSCs were mechanically dissociated and transferred in a new Matrigel-coated dish in Differentiation Medium (DM; Neurobasal medium supplemented with N2 (1X) B27 (1X), P/S (1X), NEAA (1X), Glutamax (1X) (life technologies)), ITS-A (1X, final concentration; life technologies, No. 41400045), LDN 193189 (0.5 μM, Sigma, SML0559) and CHIR99021 (3 μM, Sigma, 5ml-1046). All reagents are diluted in N2. At day 6, medium is changed with a medium containing DM, LDN 193189, IGF-1 (4 ng/ml, Peprotech, No.100-11) and HGF (10 ng/ml, Peprotech, No.100-39) and β Mercaptoethanol (ThermoFisher, No 31350010). At day 8, HGF is removed and medium is changed with DM supplemented with IGF (4ng/ml, Peprotech, No.100-11), β Mercaptoethanol. At day 12, medium is changed with DM supplemented with IGF (4ng/ml, Peprotech, No.100-11), β Mercaptoethanol and DAPT (10 μM, Tocris). Medium is changed every day until Day 17 where medium is changed with DM and IGF (4ng/ml, Peprotech, No.100-11). For ChIP-seq and RNA-seq experiments, cells were collected 30 days post differentiation.

**Transfections**

Transfections were performed with Calcium phosphate. Plasmids to be transfected are mixed with 250mM $CaCl_2$ and 2X BBS (50mM BES, 1,5mM $Na_2HPO_4$, 280mM NaCl). After 15 minutes incubation, the mixture was added dropwise directly in the culture media. Cells were incubated at 37°C and 5% $CO_2$. The media was changed 5 hours after transfection. Cells were collected 48 to 72 hrs post transfection.

**RNA extraction, quality control and library preparation**

For high-throughput RNA sequencing, total RNA was extracted using the RNAeasy kit (Qiagen) following manufacturer's instructions. Quality, quantification and sizing of total RNA was evaluated using the RNA 6000 Pico assay (Agilent Technologies Ref. 5067-1513) on an Agilent 2100 Bioanalyzer system. The RNA integrity number (RIN) was calculated for each sample and only samples with a RIN >9 were kept for further use. RNA-Seq libraries were generated from 600 ng of total RNA using TruSeq Stranded mRNA Library Prep Kit and TruSeq RNA Single Indexes kits A and B (Illumina, San Diego, CA), according to manufacturer's instructions. Briefly, following purification with poly-T oligo attached magnetic beads, the mRNA was fragmented using divalent cations at 94°C for 2 minutes. The cleaved RNA fragments were copied into first strand cDNA using reverse transcriptase and random primers. Strand specificity was achieved by replacing dTTP with dUTP during second strand cDNA synthesis using DNA Polymerase I and RNase H. Following addition of a single 'A' base and subsequent ligation of the adapter on double stranded cDNA fragments, the products were purified and enriched with PCR (30 sec at 98°C; [10 sec at 98°C, 30 sec at 60°C, 30 sec at 72°C] x 12 cycles; 5 min at 72°C) to create the cDNA library. Surplus PCR primers were further removed by purification using AMPure XP beads (Beckman-Coulter, Villepinte, France) and the final cDNA libraries were checked for quality and quantified using capillary electrophoresis. Libraries were sequenced at the IGBMC GenomeEast facility using a HiSeq 4000 1x50bp.

For quantitative PCR analysis, total RNA was extracted from cell pellet using Trizol (Thermofisher). Briefly, cell pellets were dissolved in Trizol 5:1 Chloroforme and then centrifuged to separate the different phases. Isopropanol 100% was added to precipitate the RNA overnight at -80°C. After centrifugation and washing of the RNA pellet using 70% ethanol, RNA was resuspended in ultrapure DNase-free water (Thermofisher). RNA quality was assessed by agarose gel electrophoresis and quantification using a Qubit system.

**RNA-Seq data processing and differential expression analysis.**

Fastq sequence data quality was assessed at the IGBMC facilityusing FastQC v0.11.5 and trimmed the reads to remove adapter sequences and low-quality bases using DimerRemover v0.9.2. The resulting trimmed single-end reads were aligned using STAR v2.5.3a (*6*) to the hg38 human genome release.

Reads were aligned on GRCh38 human genome reference using STAR v2.5.3a. BAM files were sorted and indexed using Sambamba (v0.6.6). Stringtie (v1.3.1c) was used to quantify aligned reads using GENCODE. Differentially expressed genes between conditions were identified using R package DESeq2 (v1.18.1). Heatmaps were realized using the R package, pheatmap (v1.0.12) using TPM counts for RNA-seq. Over representation were realized using R package clusterProfiler (v3.14.3) or Panther. DEG identified in RNA-seq with an FDR adjusted pvalue < 0.05 and abs(log2FC) > 2 were used as input. Identified GO-terms in biological process ontology were selected based on FDR adjusted pvalue < 0.05.

**Quantitative RT-PCR**

Reverse transcription of 1 µg of total RNA was performed using the Superscript IV First-Strand cDNA Synthesis kit (Life Technologies). We used a mix of oligo dT and random hexamers to target all types of RNA and followed manufacturer's instructions to synthesize cDNA. Remaining RNA was degraded by incubation with *E.coli* RNase H (Life Technologies) at 37°C for 20 minutes. Primers were designed using Primer Blast (Table S2). Real-time PCR amplification was performed on a QuantStudio 5 Real-Time PCR System (Thermofisher) using the SYBR green master mix (Roche). All PCR were performed using a standardized protocol and data were analyzed with the QuantStudio 5 Real-Time PCR System (Thermofisher). Primer efficiency was determined by absolute quantification using a standard curve. For each sample, fold-change was obtained by comparative quantification and normalization to expression of the *GAPDHD, HPRT and PPIA* housekeeping genes used as standard. Data are expressed as means ± SD.

**DNA extraction and Sodium bisulfite sequencing**

DNA was extracted from the different types of samples using the NucleoSpin Tissue (Macherey-Nagel) according to manufacturer's instructions.

For sodium bisulfite sequencing, 1 µg of genomic DNA was denatured for 30 minutes at 37°C in NaOH 0.4N and incubated overnight in a solution of 3M Sodium bisulfite pH5 and 10mM Hydroquinone using previously described protocol (Magdinier et al., 2000). Converted DNA was purified using the Wizard DNA CleanUp kit (Promega) following manufacturer's recommendation and precipitated by ethanol precipitation for 5 hours at -20°C. Primers were designed in order to amplify methylated and unmethylated DNA with the same efficiency using the MethPrimer software(Li and Dahiya, 2002) avoiding the presence of CpGs in the primer sequence (Table S3). After sequencing, sense and antisense sequences were assembled in a single sequence and bam file converted to fastq file. After trimming of each BSP primers, data were aligned using the BiQ Analyser HiMod software (http://biq-analyzer.bioinf.mpi-inf.mpg.de)(Bock et al., 2005) and processed in R (version 3.4.2). BiQ Analyzer HiMod converts sequencing data by using « 1 » for a methylated CG, « 0 » for unmethylated and « x » in case of misalignment. For each sequenced fragment, three methylation score are calculated, (i) **the CpG methylation score** of each CpG, (ii) **the sequence methylation score**, corresponding to the average methylation level of each sequence and (iii) **the global methylation score** that corresponds to the global level of methylation for each biological sample in a given region calculated as the ratio of methylated CpG with the number of aligned CpG for all sequences and CpG for a given biological sample as described in (Roche et al., 2019).

**Infinium MethylationEPIC Array**

Genome-wide DNA methylation analysis was performed by Infinium MethylationEPIC Array through Diagenode services. Genomic DNA was extracted using the NucleoSpin Tissue kit (Macherey-Nagel) from two different cell pellets for each sample. A minimum of 500ng was sent to Diagenode services for DNA methylation analysis. The analysis was mainly carried out using the R package ChAMP (Morris et al., 2014). Probes with missing values are removed, samples with more than 10% of probes with a detection p-value of greater than 0.01, probes with a detection p-value of greater than 0.01 in at least one sample and probes for which 5% of samples have a bead count less than 3 are filtered out. An annotation file is loaded that contains information about the location of probes - such as chromosome, position and nearby genes (Butcher and Beck, 2015). Probes targeting CpG sites that are near SNPs, belong to X/Y chromosomes, or align with multiple locations are filtered out. A matrix of Methylation Beta values is returned. The Beta-value indicates the percentage of copies of a CpG site from a given sample that were methylated (Du et al., 2010) determined for each CpG location as the relative intensity of methylated signal (M) and unmethylated signal (U) (Bibikova et al., 2011). Samples from the different groups were compared to identify Differentially Methylated Probes (DMPs) based on a significantly different average methylation level at CpG site. The ChAMP function for identifying DMPs uses the limma package (Ritchie et al., 2015). A 0.2 absolute difference in mean Beta values (deltaBeta) was used to identify DMPs (Smyth, 2004). The Probe Lasso method was used to identify regions of differential methylation (Differentially Methylated Regions, DMR) using the champ.DMR function. Probe Lasso starts by selecting DMPs (default p-vlaue cutoff = 0.05) and extending a lasso in either direction. The size of the lasso depends on the gene feature (Body, IGR etc) and CGI relation (Island, Shore, etc) associated with probe location. The rationale for differing lasso sizes is that probe density varies between gene features and CGI relations. Mini DMRs are created when the number of DMPs contained within a lasso is greater or equal to a minimum value. Overlapping or neighboring mini DMRs are merged until the distance between merged DMRs is at least 1,000 bp. A p-value is calculated for each DMR, using Stouffer's weighted method to combine the p-values of the probes contained in the DMR. The default cut off p-value for DMRs to be selected is 0.05, and they must contain at least 7 probes and exceed 50 bp in width. The Combat method was used to correct for batch effects (Johnson et al., 2007). Heatmaps were realized using the pheatmap (v1.0.12) R package using Beta value matrix for methylation. Violin plots were realized by converting Beta-value matrix to M-value (for better visualization) using the lumi (v2.42.0) R package then by plotting values using the ggplot2 (v3.3.3)R package. DMPs with an FDR adjusted pvalue < 0.05 and an abs(deltabeta) > 0.2 were represented as barplot using the ggplot2 (v3.3.3) R package for genomic features and CGI status according to Illumina Epic array annotation file. Karyoplots of DMPs with FDR adjsuted pvalue < 0.05 and abs(deltaBeta) > 0.2 were realized using the KaryoplteR (v1.16.0) R package. DMPs were represented as a density measured in bins. Bin size for *HOXA*,*B*,*C*,*D*, *PCDHA*,*B*,*G* = 1000bp, Bin size for Prader Willi syndrome locus and DUX4 = 10 000bp, Bin size for 4q35.1 = 20 000bp, Bin size for 4q35.2 = 50 000bp, Bin size chr6 = 1 000 000 bp. For karyoplots DMPs with FDR

< 0.05 and abs(deltaBeta) > 0.5 were plotted as dot over chromosmes with y axis ranging from 0 to 1.

**Chromatin immunoprecipitation (ChIP) and ChIP-Seq data processing.**

Between 10-20x$10^6$ cells were cross-linked with 1% formaldehyde directly in culture dishes. To this aim, the fixation solution (NaCl 0.1M, EDTA 1M, EGTA 0.5M, Tris 50mM pH8 and 1% Formaldehyde (Pierce™ 16% Formaldehyde (w/v), Methanol-free) is directly added to the culture medium and incubated for 5 minutes at RT and then 40 minutes at 4°C. Cross linking reaction was stopped by addition of Glycine (final concentration 0.125M). For ChIP-seq, we used the iDeal ChIP-seq kit for Transcription Factors and iDeal ChIP-seq kit for Histones (Diagenode). Chromatin extraction, immunoprecipitations and DNA purifications is performed as recommended by the manufacturer. For ChIP-qPCR, antibodies against SMCHD1 (Abcam ab31865 and Sigma HPA039441) were mixed at a proportion of 5μg for each per IP. ChIP-grade antibodies against CTCF, H3K4me3 and H3K27me3 (Diagenode) were used as recommended by the manufacturer. For ChIP-qPCR, primers were designed using Primer Blast and Primer3 (Table S4).

ChIP samples were purified using Agencourt AMPure XP beads (Beckman Coulter) and quantified with the Qubit (Invitrogen). ChIP-seq libraries were prepared from 10 ng of double-stranded purified DNA using the MicroPlex Library Preparation kit v2 (C05010014, Diagenode s.a., Seraing, Belgium), according to manufacturer's instructions. In the first step, the DNA was repaired and yielded molecules with blunt ends. In the next step, stem-loop adaptors with blocked 5 prime ends were ligated to the 5 prime end of the genomic DNA, leaving a nick at the 3 prime end. The adaptors cannot ligate to each other and do not have single-strand tails, avoiding non-specific background. In the final step, the 3 prime ends of the genomic DNA were extended to complete library synthesis and Illumina compatible indexes were added through a PCR amplification (7 cycles). Amplified libraries were purified and size-selected using Agencourt AMPure XP beads (Beckman Coulter) to remove unincorporated primers and other reagents. Libraries were sequenced at the IGBMC GenomEast facility using a Hiseq 4000

1x50bp. Sequence reads were mapped to reference genome hg38 using Bowtie 1.0.0 with the following parameters -m 1 --strata --best -y -S -l 40 -p 2 and delivered as BAM files along with Wig files generated using with an in-house script at IGBMC (Variable step, span=25, reads were elongated to 200b).

.BAM files were analyzed using MACS2 (v2.2.6) using input DNA as control. Peaks were identified using broad peak calling for H3K4me3 and H3K27me3 and sharp peak calling for CTCF both with a qvalue threshold < 0.05. For samples with lower chip signal (ie FSHD2 and BAMS H3K27me3) we used no model paremeter with extsize = 200bp. Peaks correspondence between replicates was assessed using MSPC to obtain consensus peaks. Using –B parameter, bedGraph files were generated and whenever possible the replicates were merged together and then converted to bigWig files for data visualization. Karyoplots of bigwig files were realized using the KaryoplteR (v1.16.0) R package. Tracks are scaled to the highest value found in the window for each mark. Peaks identified in regard to input DNA are shown as black regions above each track. Over representation were realized using the clusterProfiler (v3.14.3) R package. Genes overlapping identified peaks with at least a qvalue < 0.05 in one replicate were used as input. Identified GO-terms in biological process ontology were selected based on FDR adjusted pvalue < 0.05. Intersection of genomic coordinates from Chip-seq peaks was realized using Intervene (*18*).

We also analyzed publicly available data set for SMCHD1 distribution in HCT116 cells (https://www.ncbi.nlm.nih.gov/geo/query/acc.cgi?acc=GSM1130654). Sites with SMCHD1 enrichment were determined by peak calling and comparison between HCT116 cells and KO cells. HCT116 are XY cells and none of the peaks are present on the X chromosome, consistent with a role for SMCHD1 in the regulation of X inactivation and formation of inactive X mega domains in female cells (Wang et al., 2018). A .bed genomic coordinate file containing the coordinates of 526 peaks of SMCHD1 from a MACS2 peak calling analysis was used. From this coordinate file, a fasta .fa sequence file using bedtools getfasta was created.

**Luciferase assays.**

The different experimental DNA fragments were synthetized by GenScript or after PCR amplification and inserted into pGL3 Luciferase Reporter Vectors (pGL3 Enhancer and pGL3 Promoter, Promega) at the polylinker *Nhe*I (NEB) restriction site. Plasmids were amplified in JM109 Competent Cells (Promega) according to the manufacturer's instructions. Extraction and purification were achieved using NucleoBond® Xtra Midi / Maxi kit (Machery-Nagel). Orientation and integrity of the insert were verified by Sanger sequencing.

For each experiment, pGL3 Firefly Luciferase experimental vector has been co-transfected the with the pGL4.74[hRluc/TK] Renilla Luciferase plasmid used as transfection control. Plasmids were mixed in a 10:1 proportion respectively. For transfection, cells were plated 24 hours before transfection. Luminescence was measured using the Dual-Glo® Luciferase Assay System (Promega) kit in a GloMax® Explorer Multimode Microplate Reader. Solutions were dispensed through automatic injectors with 2 seconds delay and 10 seconds measurement for each reporter. Experiments were realized in triplicates and measures were performed in triplicates for each construct (n=9). Values were normalized to Renilla luciferase levels. Position of constructs are described in Table S5.

**Position effect variegation assays.**

pCMV derived plasmid is described in (Ottaviani et al., 2009). DNA fragments were cloned downstream of the eGFP reporter gene in pCMV vectors. DNA insert were obtained from GenScript and inserted at the *Asc*I restriction site. Details are available upon request. Sanger sequencing was performed for all selected clones containing the insert in a 5' to 3' orientation. Prior to transfection, linearization of vectors was achieved using the *Bst*XI restriction enzyme (NEB). Transfection of the linearized vectors was performed using a modified calcium phosphate method (Koering et al., 2002) optimized in order to obtain a single integration per cell (Koering et al., 2002). Three days post-transfection, the Hygromycin B selection antibiotics was added to the culture medium (Life technologies) at a final concentration of 400µg/ml. Cells were kept under permanent selection for several passages. At different time points, eGFP expression was analyzed using an Accuri flow cytometer and processed using the FlowJo software (Becton-Dickinson). The percentage of eGFP-positive cells was determined using the corresponding non-transfected cells as the baseline for autofluorescence. Mean values (M1) were used to compare fluorescence in the different samples.

**Table S1:** List of primary fibroblasts and hiPSCS clones.

BAMS-Case 1; BAMS-Case 2 and BAMS-Case 9 were described in (Gordon et al., 2017b)

All cells are described in (Badja et al., 2014; Dion et al., 2019).

| | Diagnosis | *SMCHD1* status | Age | Gender | F | iPS |
|---|---|---|---|---|---|---|
| AG08498 | Healthy | No mutation | 1 | Male | ✓ | ✓ |
| AG04148 | Healthy | No mutation | 56 | Female | ✓ | ✓ |
| 11440 | FSHD2 | c.2338+4A>G;<br>p.S754* | 37 | Male | ✓ | ✓ |
| 11491 | FSHD2 | c.5476+3 A>G;<br>r.5477_5547del;<br>p.V1826Gfs*19 | 66 | Female | ✓ | ✓ |
| 14586 | FSHD2 | c.573A>C ;<br>p.Q193P | 67 | Male | ✓ | ✓ |
| 120521C | FSHD2 | p.L1031 | 10 | Female | ✓ | ✓ |
| BAMS-1 | BAMS | c.407A>G<br>p.E136G | 5 | Male | ✓ | ✓ |
| BAMS-2 | BAMS | c.403A>T<br>p.S135C | 28 | Female | ✓ | ✓ |
| BAMS-9 | BAMS | c.1259A>T<br>p.D420V | 3 | Male | ✓ | ✓ |

**Table S2:** Sequence of the primers used for RT-qPCR. Primers for DUX4 and DUX4 targets were described in (Snider et al., 2010).

| Gene | Forward Primer | Reverse Primer |
|---|---|---|
| *HPRT* | TGATAGATCCATTCCTATGACTGTAGA | CAAGACATTCTTTCCAGTTAAAGTTG |
| *PPIA* | ATGCTGGACCCAACACAAAT | TCTTTCACTTTGCCAAACACC |
| *GAPDH* | AGCCACATCGCTCAGACAC | GCCCAATACGACCAAATCC |
| *HOTAIR* | ATGAACTGGCGAGAGGTCTG | TTCAACCCCCTCCCCCATAA |
| *HOXA11* | GGCGGCTCCAGTGGC | CGCTGAAGAAGAACTCCCGT |
| *HOXA13* | GGAACGGCCAAATGTACTGC | GATGGGAGACCACGTCGGG |
| *HOTTIP* | TTACGCCCGCAACAAAACAG | CCCTCCTTCCTTCAAACGCT |
| *HOXC4* | ATGGCCAGAGGGTTGGAAATTA | CATCCCTGAACACAGTCCGG |
| *HOXC5* | CACATGAGCCACGAGACGG | TCCACTTCATCCTGCGGTTC |
| *HOXC6* | ATGAATTCGCACAGTGGGGT | TTGATCTGTCGCTCGGTCAG |
| *HOXC8* | ACAGTAGCGAAGGACAAGGC | CCAAGGTCTGATACCGGCTG |
| *HOXC9* | CCGCAGCTACCCGGACTA | CGTGAATCCAGTTGGCCACG |
| *HOXC10* | CCCTCGGAGAGCGAAAAGG | TCAGCCAATTTCCTGTGGTGT |
| *HOXC11* | GGCTGAGGAGGAGAACACAAA | GTCGGTCCGTCAGGTTCAG |
| *HOXB2* | GAGAGACCGAAATCTCCCCC | AAGGAAGTCAGACACTCGGC |
| *HOXB3* | CCTCCCGCAAATCTCCTTGG | CCCTCCTTTGCGCCTCTC |
| *HOXB6* | GGTCTGCAAAGGCGCGG | AGGAACTCATTGGGAGGGGA |
| *HOXBAS3* | CCCTCCCTCCAAGTCCAGTA | GGATATCGCTGGGTTCCCG |
| *HOXB9* | GAGAGGCCGGATCAAACCAA | CTACGGTCCCTGGTGAGGTA |
| *HOXD1* | CTCTCTGGAACAACCCCCAC | CTGGAACTCGGAAGCCAACT |
| *HOXD3* | GTCGTCATTAATCTGCCACGC | GGCTCCAGGTGACCACAATTA |
| *HOXDAS2* | GCGATTCTTACCCGAAGGCT | AGCGCCTAGTGGATTACAGC |
| *HOXD8* | CAGTGCTGTGGTGCGAAAAT | ACACTCTGGCCTCGGTTTAC |

**Table S3:** Sequence of the primers used for Sodium bisulfite PCR. Primers for D4Z4 were described and validated in (Gaillard et al., 2014; Roche et al., 2019).

| Locus | Name | Primers | Temperature | Size |
|---|---|---|---|---|
| D4Z4 | 5P | AAATATGTAGGGAAGGGTGTAAGTT<br>CTTAAATATACCAAACCCTCTCTCC | 56°C | 341 |
| | MID | ATTTATGAAGGGGTGGAGTTT<br>ATAACCTAAACCAACCRTTCTCTA | 56°C | 419 |
| | 3P | GTTTTGTTGGAGGAGTTTTAGGA<br>CTAAACCTAAAAAACAAAAATCCCA | 56°C | 237 |
| | DR1 | GAAGGTAGGGAGGAAAAG<br>ACTCAACCTAAAAATATACAATCT | 56°C | 254 |
| HOX | HOXA13 | TGTGTTAGTTTATTTTTGGTTATGG<br>CAAACAAACCTACTTATAACTCCTC | 56°C | 216 |
| | HOXB2 | TATTTTAGATTTAATGGTGGTTGGG<br>ACCTCTAAATTTTTCATTCATTAACCTT | 56°C | 213 |
| | HOXB6 | GGGAGATAGTAAATATTTTTTTGT<br>AAAAACCAATAATTCTAACCTCC | 58°C | 269 |
| | HOXC4 | TTTTTATTTGTTTGGTTTAGTTGGG<br>CTTTTCCAAAAATCTCCATTCATAA | 56°C | 273 |

**Table S4:** Sequence of primers used for ChIP

| Gene | Forward Primer | Reverse Primer |
|---|---|---|
| BET1L | TTGCTTGCAGTGTAGTCGGG | GGGAGGAATCCCAGCAAGAT |
| CHR5 | GGAGTTGGGGAAGCTAGGAA | GATCATCCGTGGCTTGAGAT |
| D4Z4-3’ | CTCAGCGAGGAAGAATACCG | ACCGGGCCTAGACCTAGAAG |
| D4Z4-5’ | ACGACGGAGGCGTGATTT | AGTGTGGCCGGTTTGGAA |
| D4Z4-DR1 | CCCGCCTCCGGGAAAAC | GGGATGTGCGGTCTGTGAA |
| D4Z4-Mid | TCATGAAGGGGTGGAGCCTG | TCCAAACGAGTCTCCGTCGC |
| NCAM2 | CCAGACACCTCATTAGACAGCA | GTGCCCATCCTAAGCCTCTTG |
| SEMA5A | ATGTCCTAGAGGGGCAAAGC | AATGCAGACGCAAGAGCCAT |
| USP17 | TGCCTTGACATGCAGCCATA | GTGCTTTCCTGAGTGGCTCT |
| WASH7P | CGCCTTCAGAGTACCACCG | TCAGCACAGACCCGGAGA |
| DMR HOXA13 | GGCGGTGTTTACCGACTCTT | GTCGTCTGTGGAGCTGAGAC |
| DMR HOXC4/5/6 | CCCATCCAACATCCACTCCAG | CTCCATTCATGGCGATCGTG |
| DMR HOXB2 | AAACACCAGAGGACCACGAC | GCATTGGGGAGAGAGCATGT |
| DMR HOXB6 | CAGGCCCTATAGAAACCAGGAC | TTCTCTTGCCTGGTGCGGAT |

**Table S5:** Position of genomic fragment, related to the Hg38 assembly for reporter gene assays.

| Gene | Chromosome | Positions | |
|---|---|---|---|
| BET1L | Chr 11 | 190011 | 190271 |
| HOXA13 | Chr 7 | 27201926 | 27202927 |
| HOX B2 | Chr 17 | 48544650 | 48545353 |
| HOXB6 | Chr 17 | 48603749 | 48604273 |
| HOX C4 | Chr 12 | 54053160 | 54054306 |
| NCAM2 | Chr 21 | 21003843 | 21003903 |
| SEMA5A | Chr 5 | 9339379 | 9339419 |
| WASH7PL | Chr 1 | 10071 | 10271 |

### Legend to the supplementary figures

**Figure S1. Gene expression profiling in fibroblasts and induced pluripotent cells from patients carrying a mutation in *SMCHD1*. A.** Heatmap of RNAseq data (TPM values with a row sum >1, distance: Manhattan, Clustering: Ward.D2) for gene expressed in primary fibroblasts from healthy donors (controls, CT), patients affected with BAMS or FSHD2. **B.** Volcano plots for genes differentially regulated in BAMS (upper panel) or FSHD2 (lower panel) cells versus controls. Fold changes (FC log 2) are compared to the number of reads (logCounts, 58219 variables). Black dots represent genes that did not reach significance whereas dysregulated genes are shown in red. **C.** Heatmap of RNAseq data (TPM values, distance: Manhattan, Clustering: Ward.D2) for gene expressed in induced pluripotent stem cells from healthy donors (controls, CT), patients affected with BAMS or FSHD2. **D.** Volcano plots for genes differentially regulated in BAMS (upper panel) or FSHD2 (lower panel) hiPSCs versus controls. Fold changes (FC log 2) are compared to the number of reads (logCounts, 58219 variables). Black dots represent genes that did not reach significance whereas dysregulated genes are shown in red. **E.** Venn diagrams for DEGs in primary fibroblasts for the different categories of patients compared to controls and intersection with the list of DUX target genes (Geng et al., 2012). DUX4 target genes that are differentially expressed in the different categories are indicated with 6 genes in FSHD2, 14 in BAMS and 5 common to the two diseases. **F**. Venn diagrams for DEGs in hiPSCs for the different categories of patients compared to controls and intersection with the list of DUX target genes (Geng et al., 2012). Only one gene, *OAS1* is differentially expressed in BAMS cells.

**Figure S2. DNA methylation profile in control, BAMS and FSHD2 primary fibroblasts. A.** Violin plot showing the Log ratio of methylation percentage (M-values) used to derive Beta values and distribution of DNA methylation levels between samples**. B.** Density plots for normalized DNA methylation levels (Beta values) in control, BAMS and FSHD2 fibroblasts showing a bimodal distribution with peaks of unmethylated and methylated sites. A significant number of sites exhibit intermediate levels of DNA methylation. **C.** Density plots for DNA methylation levels (Beta values) after batch adjustment. **D.** Hierarchical clustering of FSHD2, BAMS and control subjects for CpGs selected among the 1000 most variable position on the basis of Beta values after stringent quality control, data normalization, and removal of probes mapping to sex chromosomes. Probes associated to the same gene were aggregated by mean.

**Figure S3. Chromosomal distribution of DMPs.** Karyoplots showing the average DMP methylation levels for the different autosomes in FSHD2 (**A,** blue) or BAMS (**B,** red) versus controls (green).

**Figure S4. DNA methylation profile of chromosome 6 in FSHD2 and BAMS fibroblasts.** DMPs were represented as a density measured in bins (bin size chr6 = 1 000 000 bp). Upper graphs correspond to hyperM DMP, lower graph to hypoM DMPs in FSHD2-11440, FSHD2-11490 (**A**), BAMS1 and BAMS9 (**B**) primary fibroblasts. Differential methylation is determined in comparison to healthy controls.

**Figure S5. DNA methylation profile in control, BAMS and FSHD2 induced pluripotent stem cells. A.** Violin plot showing the Log ratio of methylation percentage (M-values) used to derive Beta values and distribution of DNA methylation levels between hiPSC samples**. B.** Density plots for normalized DNA methylation levels (Beta values) in control, BAMS and FSHD2 hiPSCs showing a bimodal distribution with peaks of unmethylated and unmethylated sites. A significant number of sites exhibit intermediate levels of DNA methylation. **C.** Cumulative histograms of the distribution of hypo- and hypermethylated probes in BAMS and FSHD2 induced pluripotent stem cells. **D.** Cumulative histograms for DMP by CpG content relative to CpG islands, shores (2 kb flanking CpG islands), shelves (2 kb extending from shores) or open seas (isolated CpG in the rest of the genome) in BAMS and FSHD2 hiPSCs. **E.** Cumulative histograms for DMP by features corresponding to genes first exon, 3’ UTR,

5’UTR, gene bodies, Exon boundaries, Internal genomic regions (IGR), probes located 1500 bp from transcription start sites (TSS1500) or 2000 bp from transcription start sites (TSS200), in BAMS and FSHD2 hiPSCs. Values are presented in supplementary table 3. **F.** Venn Diagram for DMRs in BAMS and FSHD2 hiPSCs.

**Figure S6. DNA methylation profiling of *HOXB* and *D* genes in SMCHD1-deficient primary fibroblasts.**

**A.** Representative distribution of the DNA methylation profile of DMR at the *HOXB* locus in the different FSHD2 (11440, 11491, 14586, red) or BAMS (BAMS1, BAMS2, BAMS9, Cyan) samples. **B**. Hierarchical clustering of FSHD2, BAMS and control subjects based on Beta values of aggregated probes (mean) to HOXB genes (chr17: 48,496,690-48,772,940, distance: Canberra, clustering Ward D2). **C**. Representative distribution of the DNA methylation profile of DMR at the *HOXD* locus in the different FSHD2 (11440, 11491, 14586, red) or BAMS (BAMS1, BAMS2, BAMS9, Cyan) samples. **D**. Hierarchical clustering of FSHD2, BAMS and control subjects based on Beta values of aggregated probes (mean) to HOXD genes (chr2: 176,069,968-176,201,505 (Gordon et al., 2017a; Lemmers et al., 2012; Shaw et al., 2017), distance: Canberra, clustering Ward D2).

**Figure S7. Bisulfite sequencing for DNA methylation profiling of *HOX* genes.** DNA methylation was determined after sodium bisulfite modification for 4 *HOX* genes DMRs (*HOXA13*, *HOXB2*, *HOXB6* and *HOXC4/5/6*) for primary fibroblasts (**A**) or corresponding hiPSC clones (**B**). For each sequence, cumulative histogram bars represent the percentage of methylated CpG at each position of the sequence analyzed (black) and the percentage of unmethylated CGs (white bars).

**Figure S8. Analysis of *HOX* genes in wild-type HEK and HEK *SMCHD1*-KO cells.**

**A.** We analyzed SMCHD1 binding to the different HOX DMRs (*HOXA13*, *HOXB2*, *HOXB6* and *HOXC4/5/6) in* HEK *and* HEK-*SMCHD1*-KO cells. Enrichment over input was determined by qPCR. Bar plots display the average enrichment after normalization over a single copy intergenic region. Values are average from at least three independent biological replicates and a technical duplicate for each. Error bar represents standard error. Statistical significance was determined using a paired two-tailed student's t test (* $p< 0.01$; ** $p< 0.001$). **B.** Cumulative histogram display DNA methylation profiles analyzed in HEK cells and cells in which *SMCHD1* gene expression was invalidated by gene editing (HEK KO) for *HOXA13*, *HOXB2*, *HOXB6* and *HOXC4/5/6* DMRs. Black bars correspond to the percentage of methylated sites, white bars to unmethylated sites. **C.** Expression of different *HOX* genes was evaluated by RT-qPCR performed as biological and technical in triplicates in HEK and HEK KO cells. Expression was normalized to three housekeeping genes (*GAPDH*, *HPRT* and *PPIA*). Statistical significance was determined by Kruskal-Wallis statistical test. ** p-value<0.005, **** p-value<0.00005.

**Figure S9. Somatic loss of heterozygosity at the 18p11.32 locus encompassing the *SMCHD1* gene is not associated with D4Z4 hypomethlation in breast cancer tumors.** We analyzed the methylation level of the different regions within D4Z4 (DR1, 5P, MID and 3P) in 29 breast cancer tumors with a loss of 18p heterozygosity encompassing the *SMCHD1* gene locus (**A**). **B**. Graphs display the methylation rate (*y-axis*) related to the log2 ratio (*x-axis*) of 18p loss in the different samples as determined by comparative genomic hybridization (CGH). The correlation value and level of significance is indicated for the different samples.

**Figure S10. Chromatin profiling in BAMS and FSHD2 muscle cells.**

**A**. Distribution of CTCF, H3K4me3 and H3K27me3 relative to the different autosomes in Controls (CT2, black bars), BAMS (cyan bars) or FSHD2 (red bars) cells. **B**. Distribution of peaks width for CTCF, H3K4me3 or H3K27me3 in Control BAMS1 and FSHD2 muscle fibers. Statistical significance was determined using a OneWay ANOVA with Brown-Forsythe multiple comparison, **** p-value<0.0001.

**Figure S11. Chromatin profiling of *HOXB* and *D* genes in *SMCHD1*-deficient cells.**

**A**. Profiling of the *HOXB* genes locus (*HOXB* : chr17: 48,520,200-48,740,600 ; 220400 bp) in the different samples (Controls, black; BAMS, Cyan or FSHD2, red) for H3K4me3, H3K27me3 or CTCF enrichment. **B**. Profiling of the *HOXD* genes locus (chr2: 176,090,006-176,190,745 ; 100739 bp) in the different samples (Controls, black; BAMS, Cyan or FSHD2, red) for H3K4me3, H3K27me3 or CTCF enrichment. **C**. Venn diagram of HOX genes differentially expressed in hiPSC-derived muscle cells. Fold change (FC) for the different genes and different categories are indicated next to each group. Red dots indicate genes that are upregulated ; green dots, downregulated ones.

**Figure S12. For a number of sites, loss of SMCHD1 binding leads to increased CTCF enrichment.**

Relative quantification of SMCHD1 or CTCF enrichment after ChIP-qPCR in HEK293 cells and HEK293 KO cells for putative SMCHD1 sites selected from publicly available ChIP-Seq data (https://www.ncbi.nlm.nih.gov/geo/query/acc.cgi?acc=GSM1130654) at the *NCAM2*, *BET1L*, *SEMA5A* and *WASH7P* loci. The different proteins targeted by immunoprecipitation are indicated on the *x-axis* (SMCHD1, CTCF). Each experiment was done in two biological replicates with each biological replicate analyzed in technical duplicates. CT values were normalized to the input. Fold change was calculated and values were normalized to an intergenic unique sequence on chromosome 5. Statistical significance was determined using a 2way ANOVA test for multiple comparisons (* p value < 0.1; **** p value < 0.001).

**Figure S13. D4Z4 subfragment harbor SMCHD1-dependent *cis*-regulatory activity**

**A.** Schematic representation of the D4Z4 element from position 1 to 3303 given relative to the two flanking *Kpn*I sites (K) (to scale). The different regions within *D4Z4* are indicated: *LSau* repeat (position 1-340), Region A (position 869-1071), *hhspm3* (position 1313-1780), *DUX4* ORF (position 1792-3303). Different fragments in the proximal part of the repeat encompassing to the most differentially methylated regions (DR1, position 566-819 and 5P, position 1027-1253 relative to the first *Kpn*I site) were cloned in the pGL3 basic vector lacking the SV40 promoter and enhancer, the pGL3 Enhancer vector lacking the SV40 promoter or the pGL3 Promoter vector lacking the SV40 enhancer. Expression of the firefly luciferase was determined 48 hrs post-transfection in HEK cells or HEK KO cells. Firely luciferase levels were normalized to expression of the Renilla luciferase used as transfection control. Values corresponding to the luciferase activity (expressed in arbitrary units, A.U.) are the average of at least three independent assays with three measures per experiment (n=9). Error bar represents standard error. Statistical significance was determined using a Mann-Whitney test, * p value = 0.1; ** p value <0.01, *** p value < 0.001; **** p value < 0.0001. **B.** Relative luminescence activity (RLU) for the different constructs indicated on the left of the histogram after transfection in HEK (black bars) or HEK-KO cells (grey bars). **C**. Different subfragments corresponding to the DR1 fragment were cloned in the pGL3 promoter vector. This sequence contains 31 CpG sites that are differentially methylated in cells from patients with a mutation in SMCHD1 (BAMS and FSHD). Fragments containing CG1-10; 10-20 or 21-30 were tested together with fragments lacking these different elements (ΔCG1-20; ΔCG10-20; ΔCG21-31). **D**. Different sequences to be tested were cloned downstream of the *eGFP* reporter gene: *HOXB2* DMR, *HOXB6* DMR, *SEMA5A*, *NCAM2*, *BET1L, WASH7PL*. Linearized plasmids were transfected into cells. Stable eGFP expression was measured by flow cytometry (FACS) for an extended period of time in cells grown in the presence of Hygromycin B. Representative spectra of the % of eGFP positive cells a presented. eGFP expression level is the average of 5 measurements from day 18 to day 40 post-transfection, when *eGFP* expression reaches a plateau, of three independent assays ± S.D. For each condition, eGFP expression was compared to values obtained in cells transfected with the empty vector (pCMV). In HEK KO cells, eGFP expression was also measured 72 hrs after transfection of a SMCHD1 expression vector (grey curves). Asteriks indicate statistically significant values relative to control vectors (pCMV) (Student's t test). * $p < 0.001$; ** $p < 0.005$; *** $p < 0.05$.
